# Loop Catalog: a comprehensive HiChIP database of human and mouse samples

**DOI:** 10.1101/2024.04.26.591349

**Authors:** Joaquin Reyna, Kyra Fetter, Romeo Ignacio, Cemil Can Ali Marandi, Astoria Ma, Nikhil Rao, Zichen Jiang, Daniela Salgado Figueroa, Sourya Bhattacharyya, Ferhat Ay

## Abstract

HiChIP enables cost-effective and high-resolution profiling of chromatin loops. To leverage the increasing number of HiChIP datasets, we developed Loop Catalog (https://loopcatalog.lji.org), a web-based database featuring loop calls from 1000+ distinct human and mouse HiChIP samples from 152 studies plus 44 high-resolution Hi-C samples. We demonstrate its utility for interpreting GWAS and eQTL variants through SNP-to-gene linking, identifying enriched sequence motifs and motif pairs, and generating regulatory networks and 2D representations of chromatin structure. Our catalog spans over 4.19M unique loops, and with embedded analysis modules, constitutes an important resource for the field.

## BACKGROUND

Changes in chromatin folding caused by mutations and variants can impact cell-type-specific function and disease risk via altered 3D interactions between genetic loci (Chakraborty & Ay, 2019; S. Wang et al., 2023). An important feature of chromatin folding is the existence of chromatin loops that connect regions separated by large genomic distances, often up to a megabase but with notable cases spanning even larger distances (Chandra et al., 2021; Friman et al., 2023; Levo et al., 2022; Long et al., 2020). These loops can be broadly categorized into: *i)* structural loops demarcating domains of interactions such as topological domains (TADs) and marked by the binding of CTCF and cohesin at their anchors; *ii)* regulatory loops which join distal gene regulatory elements such as enhancers and promoters to modulate gene expression (Boltsis et al., 2021; Brandão et al., 2021; L.-F. Chen & Long, 2023; Hu et al., 2023; Ibrahim & Mundlos, 2020; Ito et al., 2022; Popay & Dixon, 2022; Portillo-Ledesma et al., 2023; Qiu et al., 2023; Schaeffer & Nollmann, 2023; Xu et al., 2020). Our understanding of the exact role that these loops/interactions play in cell-type specific gene regulation and ultimately disease susceptibility is far from complete.

To capture such interactions, the Hi-C procedure was developed as a high-throughput genome-wide assay that carries proximity ligation either in dilution (Lieberman-Aiden et al., 2009) or in-situ within intact nuclei (S. S. P. Rao et al., 2014). As a subset of interactions, regulatory loops are often associated with regions enriched for histone modifications such as H3K27ac and H3K4me3, transcription factors, or accessible chromatin (Davies et al., 2016; Fang et al., 2016; Fullwood et al., 2009; S. Liu et al., 2022; Mumbach et al., 2016; Wei et al., 2022). The development of the HiChIP (Hi-C with chromatin immunoprecipitation) assay represents an extension of the in-situ Hi-C methodology that can facilitate the capture of these protein-specific regulatory loops by using immunoprecipitation to enrich for interactions involving anchors associated with the binding of transcription factors (TFs) or histone modifications of interest. Given the ability to enrich signal for a subset of the genome (e.g., active regulatory elements), HiChIP enables high-resolution profiling of chromatin interactions of interest with lower sequencing depth compared to Hi-C (Fang et al., 2016; Mumbach et al., 2016; Yu et al., 2021). Another advantage of HiChIP, at the time of its initial development, was its lower cell input requirement (1-5M), hence enabling the characterization of 3D organization in primary cells. Since the introduction of HiChIP in 2016, the number of publicly available human and mouse HiChIP studies published has consistently increased (**Figure 1A**). This indicates the growing popularity of the HiChIP assay, particularly as a method of investigating distal interactions between genetic variants, often in non-coding regions of the genome, and their potential target genes in a cell-type and context-specific manner (Chandra et al., 2021; Duan et al., 2021; Giambartolomei et al., 2021; Mumbach et al., 2017; O’Mara et al., 2019; Shi et al., 2021; W. Wang et al., 2021).

**Figure 1.**
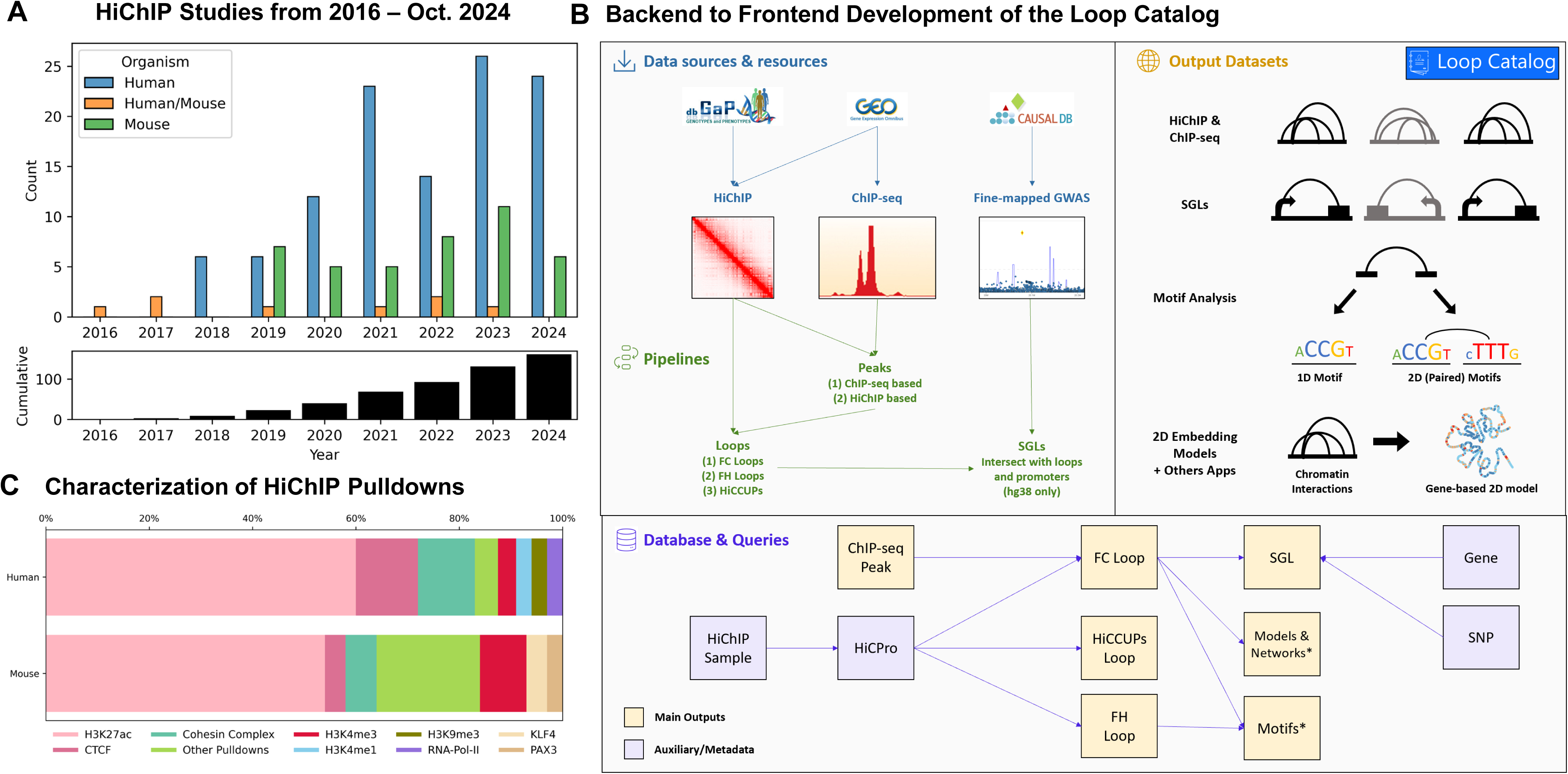
High-level summary of the Loop Catalog. **A)** Breakdown of HiChIP Samples from 2016 to 2024. The top panel shows the number of studies broken down by human (blue), mouse (teal), or both (orange). Bottom panel shows a cumulative breakdown. **B)** Schema for the development of the Loop Catalog starting from raw sequencing files to processing (top left), database storage (bottom) and web accessible analyses (top right). **C)** Breakdown of samples in the Loop Catalog by target protein or histone modification and organism.

In the context of human disease, HiChIP and similar assays provide a 3D view for the annotation of disease associations of non-coding genetic variants identified from genome wide association studies (GWAS) (Fu et al., 2018; Gazal et al., 2022; Giambartolomei et al., 2021; Orozco et al., 2022; Zhong et al., 2022). Combined with efforts from multiple large consortia for cataloging putative regulatory elements spanning distinct cell types (e.g., ENCODE, BLUEPRINT and Roadmap Epigenomics), mapping such 3D maps of chromatin organization has become a critical piece of the puzzle, which led to formation of the 4D Nucleome Consortium (Dekker et al., 2017; Reiff et al., 2022). In parallel, other large-scale efforts identified SNPs (single nucleotide polymorphisms) associated with gene expression (i.e., expression quantitative trait loci or eQTLs) for different tissues and primary cell types (Bernstein et al., 2010; ENCODE Project Consortium et al., 2022; GTEx Consortium, 2020; Martens & Stunnenberg, 2013; Ota et al., 2021; Schmiedel et al., 2021). Although multiple studies incorporated GWAS, eQTL, and high-resolution chromatin looping across many different matched/related cell types, it still remains a challenge to find a catalog of uniformly processed loops identified from datasets generated outside the ENCyclopedia Of DNA Elements (ENCODE) and 4D Nucleome (4DN) consortia. Recent publications and databases, namely HiChIPdb (Zeng et al., 2023), ChromLoops (Zhou et al., 2023), Cohesin-DB (J. Wang & Nakato, 2023) and 3DIV (Jung et al., 2019; Kim et al., 2021), were first attempts to catalog loops by curating them from the broader literature. We provide a detailed comparison of the Loop Catalog with these previous databases on the basis of the number of samples processed, the choice of tools for data processing including loop calling, integrated visualization and additional data types compiled (**Table 1**). In addition to possessing the largest number of HiChIP samples, unique features of the Loop Catalog include utilization of matched chromatin immunoprecipitation-sequencing (ChIP-seq) data when available, enabling a broader set of data download and visualization capabilities, and additional modalities including SNP-to-gene linking, traditional and pairwise motif enrichment analysis, regulatory network analysis, and 2D representations of chromatin structure (**Table 1**).

**Table 1.** Comparison of processing methods and website features of Loop Catalog, ChromLoops, HiChIPdb, Cohesin-DB, and 3DIV. Categories include specification and quantification of data types, implementation of loop calling (software, configurations, replicates), ease of data download and visualization, including the ability to select and visualize multiple samples simultaneously, and embedded biological analysis modules.* annotation of genes, SNPs, E/Ps, silencers, circRNA, TWAS, chromosome open access data, alternative splicing, transcription factors ** annotation of genes, SNPs *** annotation of genes

| General Feature | Specific Feature | Loop Catalog | ChromLoops<br>(Zhou et al., 2023) | HiChIPdb<br>(Zeng et al., 2023) | Cohesin-DB<br>(Wang et al., 2022) | 3DIV<br>(Kim et al., 2021) |
| --- | --- | --- | --- | --- | --- | --- |
| Data Types | Organism | Homo sapiens, Mus musculus | Homo sapiens, Mus musculus, 11 others | Homo sapiens | Homo sapiens | Homo sapiens |
|  | Reference Genome | hg38, mm10 | hg38, mm10, 11 others | hg19 | hg38 | hg19 |
|  | Major Assays of Interest | HiChIP (n=1319, 1031 distinct)<br>Hi-C (n=44) | HiChIP (n=772)<br>PLAC-seq (n=89)<br>ChIA-PET (n=789) | HiChIP (n=200) | HiChIP (n=42)<br>Hi-C (n=385)<br>ChIA-PET (n=119) | PCHi-C (n=28)<br>Hi-C (n=401) |
| HiChIP Processing | Loop Calling | FitHiChIP, HiCCUPS | ChIA-PET Tool (V3) | FitHiChIP, hichipper | HiCCUPS | N/A |
|  | Loop Resolutions | 5kb, 10kb, 25kb | variable | 1kb, 5kb, 10kb, 50kb, variable | 5kb, 10kb, 25kb | N/A |
|  | Peak Type Used for Loop Calling | HiChIP-Inferred, ChIP-seq when available | HiChIP-Inferred | HiChIP-Inferred | N/A | N/A |
|  | Replicate Handling | Technical/Biological Reps Merged, Multiple Donors Merged | Biological Reps Merged | Technical/Biological Reps Merged | Technical Reps Merged | N/A |
|  | Pipeline Code Released | ☑ | ☑ | ☐ | ☑ | N/A |
| Data Download | Loop Calls | ☑ | ☑ | ☑ | ☑ | ☑ |
|  | Peak Calls Used for Loop Calling | ☑ | ☐ | ☐ | ☐ | N/A |
|  | Browser Track Files | ☑ | ☐ | ☐ | ☐ | ☐ |
| Data Visualization | Embedded Visualization | Epigenome Browser (WashU) | Epigenome Browser (WashU) | IGV | Epigenome Browser (WashU) | Custom |
|  | Multi-Sample Selection | ☑ | ☐ | ☑ | ☑ | ☑ |
|  | Multi-Sample Visualization | ☑ | ☐ | ☐ | ☑ | ☑ |
| Data Analysis | Synopsis | SGL Analysis, Motif Enrichment Analysis for Conserved Anchors, Loop Motif Pair Analysis, 2D Embeddings, Community Structure Analysis | Embedded Tools & Functional Anchor Annotation*, Cancer High-Frequency Loops, Species-Specific High Frequency Loops | Functional Anchor Annotation** | Functional Anchor Annotation***, Loop discovery given genome regions, Prediction of regulatory sites and target genes | N/A |

Overall, the Loop Catalog developed here is a database containing the largest set of uniformly processed HiChIP data to date, curated from over 150 publications and a total of 1334 samples (from 2016 to January 2024) leading to over 4.19 million unique looping interactions at 5 kb resolution (approximately 15.5M total across all resolutions with 11M for human, 4.5M for mouse) with an accompanying web server that enables visualization, querying and bulk download of looping data (**Figure 1B, Supp. Figure S1**). In addition to allowing easy access to this comprehensive catalog of chromatin looping data, we also demonstrated three different applications of how this data can be utilized. In the first application, we intersected fine-mapped GWAS SNPs from CAUSALdb for four autoimmune diseases (J. Wang et al., 2020) with HiChIP loops from various immune cell types to identify potential target genes for these disease-associated SNPs in each cell type. More specifically, we located loops where a SNP and a gene promoter overlapped opposing anchors of the loop and we termed these constructs SNP-to-gene links (SGLs). Across all diseases we located 3048 unique SNPs, 1486 genes, 3411 loops and 13672 SGLs that span a median genomic distance of ~140kb. One example was a fine-mapped Type 1 Diabetes (T1D) GWAS SNPs which was linked to multiple genes through HiChIP loops in multiple lymphoid cells but not in monocytes. As the second application, we investigated TF motifs in loop anchors using motif enrichment analysis (Bailey et al., 2015) and pairs of TF motifs at loop anchors through pairwise motif enrichment using a bootstrapping approach. Our analysis of regulatory loop anchors conserved across a large majority of samples identified hundreds of enriched motifs including known and novel zinc finger transcription factors some of which were also enriched for significantly overrepresented combinations of motif pairs across samples. In addition to visualization through Epigenome Browser, we also provided a network visualization for loops to highlight connected regulatory elements including enhancers and promoters. Topological structures such as communities and subcommunities in these networks alongside measurements of their connectivity properties can be explored further through an interactive user interface. For the last application, we utilized METALoci to generate 2D embedding models for visualization of chromatin conformation data together with 1D signals (e.g., ChIP-seq). This approach enables assessment of spatial autocorrelation of overlaid 1D signals using metrics established in geographical information systems (e.g., Moran’s I index). The large set of loops cataloged in this work will enable not only 3D-informed prioritization of genetic variants and enhancers involved in gene regulation but also will stimulate development of machine learning, deep learning, and network construction approaches that require large scale data.

## CONSTRUCTION AND CONTENT

### Curating HiChIP and ChIP-seq Samples

To identify a comprehensive list of publicly-released HiChIP datasets, we developed a pipeline that scans NCBI’s Gene Expression Omnibus (GEO) database (Barrett et al., 2013) for studies performing HiChIP experiments. To extract information on these studies, the BioPython.Entrez (Buchmann & Holmes, 2019) and metapub.convert (https://pypi.org/project/metapub/) packages were used. Raw sequencing data associated to these studies was then identified from the Sequence Read Archive (SRA) database using the *pysradb* Python package (https://github.com/saketkc/pysradb), and the results were manually examined to extract HiChIP samples. ChIP-seq samples corresponding to these studies were also extracted if there was a record of them within the same GEO ID as the HiChIP sample.

### Populating Sample Metadata

To automatically add metadata such as organ and cell type to the curated HiChIP data, the BioPython.Entrez package was used to perform a search query within the GEO database. GSM IDs from the previous query were used with an esummary query to the biosample database (Buchmann & Holmes, 2019). From these results, organism, biomaterial, celltype, GSM ID, SRA ID, disease, organ, treatment, tissue, and strain (only for mouse) were extracted. To improve classes for organ and biomaterial, a dictionary of classes as keys and synonyms as value (e.g. heart is a key and its synonyms are cardiovascular, atrium, aorta, etc.) was used to search across GEO and Biosample reports. “N/A” was used to indicate fields that were not applicable for a given sample, and, for the remaining fields, “Undetermined” was assigned when the information could not be retrieved.

### Downloading Raw Sequencing Datasets

HiChIP FASTQ files were systematically downloaded using one of three methods and prioritized in the following order: SRA-Toolkit’s prefetch and fasterq-dump (2.11.2) (https://hpc.nih.gov/apps/sratoolkit.html), grabseqs (0.7.0) (Taylor et al., 2020), or European Bioinformatics Institute (EBI) URLs generated from SRA Explorer (1.0) (Ewels, 2024). In addition, FASTQ files for phs001703v3p1 and phs001703v4p1 which pertain to two previously published HiChIP studies from the Database of Genotypes and Phenotypes (dbGaP) were downloaded (Schmiedel et al., 2021; Chandra et al., 2021; Sayers et al., 2022) (**Supp. Table S1**). Additionally, ChIP-seq FASTQ files were downloaded from GEO using grabseqs (0.7.0).

### Processing Reads from HiChIP Data

HiChIP sequencing reads from human samples were aligned to hg38 while reads from mouse samples were aligned to mm10 using the HiC-Pro (3.1.0) pipeline (Servant et al., 2015) (**Supp. Table S2**). Reference genomes were downloaded for hg38 (https://hgdownload.soe.ucsc.edu/goldenPath/hg38/bigZips/) and for mm10 (https://hgdownload.soe.ucsc.edu/goldenPath/mm10/bigZips/) and indexed with bowtie2 (2.4.5) (Langmead et al., 2019) using default parameters. Chromosome size files for hg38 and mm10 were additionally retrieved from these repositories. FASTQ files for all technical replicates of a HiChIP biological replicate were processed together by HiC-Pro. Restriction enzymes and ligation sequences were determined during the HiChIP literature search (**Supp. Table S3**), and restriction fragments were generated with the HiC-Pro *digestion.py* tool using default parameters. FASTQ files were split into chunks of 50,000,000 reads using the HiC-Pro *split_reads.py* utility, and HiC-Pro was run in parallel mode with a minimum mapping quality (MAPQ) threshold of 30. Invalid pairs (singleton, multi-mapped, duplicate, etc.) were removed. Raw and ICE (Iterative Correction and Eigenvector Decomposition) normalized contact matrices were generated at the 1kb, 2kb, 5kb, 10kb, 20kb, 40kb, 100kb, 500kb, and 1mb resolutions. For all other HiC-Pro configuration parameters, the defaults were used. Exceptions to this workflow are as follows:

1. For MNase HiChIP samples, R1 and R2 FASTQ reads were trimmed from the 3’ end to 50bp to account for the increased likelihood of chimeric reads. HiC-Pro was run with MIN_CIS_DIST=1000 specified to discard non-informative pairs. Validpairs generated from this HiC-Pro configuration were used for loop calling. HiC-Pro was additionally run for all MNase HiChIP samples without the MIN_CIS_DIST parameter specified to enable the collection of short validpairs which are required by the FitHiChIP *PeakInferHiChIP.sh* utility (10.0) (Bhattacharyya et al., 2019) to infer peaks from HiChIP. Validpairs generated from this HiC-Pro configuration were used only for peak inference from HiChIP data.
2. For Arima HiChIP samples originating from studies dating post-late-2022 and with an R1 or R2 mapping percentage less than 80% reported by our default HiC-Pro configuration, 5bp were trimmed from the 5’ end of the R1 and R2 FASTQ reads to remove the Arima adaptor sequence. HiC-Pro was then re-run as initially specified.

Juicer (2.0; juicer_tools.jar 2.20.00) (Durand et al., 2016) was applied to a subset of HiChIP samples. Reference genomes for hg38 and mm10 were obtained as previously described and indexed using BWA *index* (0.7.17) (H. Li, 2013) with default parameters. Restriction enzymes were determined as previously described, and restriction sites were generated using the Juicer *generate_site_positions.py* utility. HiChIP technical replicates were combined and processed together. *juicer.sh* was run with the appropriate indexed reference genome, chromosome sizes file, restriction enzyme, and restriction sites. HiChIP FASTQs were split into chunks of 50,000,000 reads (-C 50000000) and the --assembly flag was specified to generate the old style merged_nodups pairs file. For MNase HiChIP samples, the restriction enzyme was set to “none” (-s none) and no restriction site file was provided. Pairs were filtered according to a minimum MAPQ threshold of 30. Statistics reported in the inter_30.txt file were extracted for alignment tool comparison analyses.

The distiller-nf pipeline (0.3.4) (Open2C et al., 2024) was run using Nextflow (22.10.7) (Di Tommaso et al., 2017) for a subset of HiChIP samples. Reference genomes for hg38 and mm10 were downloaded and indexed with BWA as previously described. HiChIP technical replicates were combined and processed together. distiller-nf was run with the appropriate reference genome and chromosome sizes file. Mapping was performed with a chunk size of 50,000,000 reads and with the use_custom_split option set to “true”. Read alignments were parsed with options --add-columns mapq and --walks-policy mask. Duplicates were detected with max_mismatch_bp set to 1. Pairs with the UU (unique-unique) or UR/RU (unique-rescued) classification and with a MAPQ of at least 30 were retained and used for downstream analyses.

### Processing Reads and Calling Peaks for ChIP-seq Data

We utilized our previously developed ChIPLine pipeline (https://github.com/ay-lab/ChIPLine) for ChIP-seq processing (**Supp. Table S4**). ChIP-seq reads were aligned to h38 for human samples or to mm10 for mouse samples with bowtie2 (2.4.5) with the -k 4 and -X 2000 options specified and defaults otherwise. Uniquely mapped reads were determined using samtools (1.9) (Danecek et al., 2021) to remove random and mitochondrial reads and to impose a minimum MAPQ threshold of 30. PCR duplicates were marked and removed using Picard Tools (2.7.1) (*Picard Toolkit*, 2019) with a validation stringency of LENIENT. Several data quality metrics were computed through cross-correlation analysis using the *run_spp.R* utility from phantompeakqualtools (1.14) (Kharchenko et al., 2008; Landt et al., 2012). Alignment files were converted into bigwig format for visualization using (1) the UCSC *fetchChromSizes*, *genomeCoverageBed*, *bedSort*, and *bedGraphToBigWig* utilities (Kent et al., 2010) for generation of unnormalized bigwigs and (2) deepTools *bamCoverage* (3.5.1) (Ramírez et al., 2016) with -of bigwig, -bs 10, --normalizeUsing RPKM, and -e 200 options specified for generation of bigwigs normalized with respect to coverage. Narrow peaks were derived using MACS2 (2.2.7.1) (Zhang et al., 2008) (**Supp. Table S5**) with the --nomodel, --nolambda, --shift 0, --extsize 200, and -p 1e-3 parameters specified. For ChIPMentation data, a shifted tagAlign file for peak calling was first constructed by shifting the forward strand by 4bp and reverse strand by 5bp to cover the length of the transposon, and the --bw 200 parameter was additionally specified for MACS2. If available, the corresponding input ChIP-seq files were used by MACS2 for peak calling using the input vs. treatment mode. MACS2 peak calls were made without an input file if no such file was available for the sample. For ChIP-seq datasets with multiple biological replicates, the sequence files for all replicates were processed together to obtain the set of pooled ChIP-seq peaks. Narrow peaks were filtered using a Q-value threshold of 0.01 before use in all downstream analyses.

### Calling Peaks from Reads of HiChIP Data

In the absence of corresponding ChIP-seq data for a given HiChIP sample from the same study, we inferred peaks directly from HiChIP. We called HiChIP peaks with the *PeakInferHiChIP.sh* utility script from FitHiChIP, which requires the reference genome, read length (as reported by SRA Run Selector), and processed interaction pairs generated by HiC-Pro (valid, religation, self-circle, and dangling-end pairs) as input and utilizes MACS2 for peak calling (**Supp. Table S5**). If read lengths differed across technical replicates for a single HiChIP biological replicate, the mode read length was used. For cases in which there was no single mode read length, the longest read length was selected. For merged biological replicates, the mode read length of all individual biological replicates for a given sample was chosen, and similarly, the longest read length was selected in the case of multiple modes.

For comparison purposes, for a subset of HiChIP samples, HiChIP-Peaks *peak_call* (0.1.2) (Shi et al., 2021) was run using the correct chromosome size file and HiC-Pro interaction pairs. Restriction fragments were generated with the HiC-Pro *digestion.py tool* using default parameters. We used a false discovery rate (FDR) of 0.01 (-f 0.01) for peak calling with HiChIP-Peaks.

### Performing Recall Analysis of 1D Peaks Inferred from HiChIP Data

Peaks called from ChIP-seq datasets were considered the ground truth peak set. Peaks inferred directly from HiChIP datasets were assessed for their overlap with ChIP-seq peaks (**Supp. Figure S2**), and we computed the percentage of ChIP-seq peaks recovered using HiChIP-inferred peaks. HiChIP and ChIP-seq peak sets were intersected using bedtools *intersect* (2.30.0) (Quinlan & Hall, 2010) allowing for 1kb slack on both sides of the peak.

### Integration of Biological Replicates of HiChIP Data

In order to generate more deeply sequenced contact maps from the initial set of samples, we combined HiChIP biological replicates from the same study for both human and mouse datasets (**Supp. Table S1**). Before merging biological replicates, reproducibility was assessed using hicreppy (0.1.0) which generates a stratum-adjusted correlation coefficient (SCC) as a measure of similarity for a pair of contact maps (https://github.com/cmdoret/hicreppy) (**Supp. Table S6**). Briefly, contact maps in hic format were converted to cool format for hicreppy input at 1kb, 5kb, 10kb, 25kb, 50kb, 100kb, 250kb, 500kb, and 1mb resolutions using *hic2cool* (0.8.3) (https://github.com/4dn-dcic/hic2cool). *hicreppy* was run on all pairwise combinations of biological replicates from a given HiChIP experiment. First, hicreppy *htrain* was used to determine the optimal smoothing parameter value *h* for a pair of input HiChIP contact matrices at 5kb resolution. *htrain* was run on a subset of chromosomes (chr1, chr10, chr17, and chr19) and with a maximum possible h-value of 25. Default settings were used otherwise. Next, hicreppy *scc* was run to generate a SCC for the matrix pair using the optimal *h* reported by *htrain* and at 5kb resolution considering chr1, chr10, chr17, and chr19 only. A group of HiChIP biological replicates were merged if all pairwise combinations of replicates in that group possessed a SCC greater than 0.8 (**Supp. Figure S3**). We merged biological replicates by concatenating HiC-Pro valid pairs files before performing downstream processing.

### Identifying Significant Chromatin Loops from HiChIP Data

Loop calling was performed for both unmerged and merged HiChIP biological replicates by (a) HiCCUPS (JuicerTools 1.22.01) (S. S. P. Rao et al., 2014), (b) FitHiChIP with HiChIP-inferred peaks (FH loops), and (c) FitHiChIP with ChIP-seq peaks (FC loops), when available (Durand et al., 2016) (**Supp. Table S7**). Briefly, HiC-Pro validpairs files for each sample were converted to .hic format using HiC-Pro’s *hicpro2juicebox* utility with default parameters. HiCCUPS loop calling was initially performed for chr1 only using vanilla coverage (VC) normalization with the following parameters: --cpu, --ignore-sparsity, -c chr1, -r 5000,10000,25000, and -k VC. Samples which passed the thresholds of at least 200 loops from chromosome 1 for human samples and at least 100 loops from chromosome 1 for mouse samples, both at 10kb resolution, were processed further with HiCCUPS for genome-wide loop calling using the same parameters. For comparison purposes, for a subset of samples, HiCCUPS loop calling was performed with alternative normalization approaches: KR (Knight Ruiz), VC_SQRT (square root vanilla coverage), and SCALE by specifying the -k KR, -k VC_SQRT, and -k SCALE parameters respectively. FitHiChIP peak-to-all loop calling was run with HiC-Pro validpairs as input at the 5kb, 10kb, and 25kb resolutions for both the Stringent (S) and Loose (L) background models and with coverage bias regression, merge filtering, and an FDR threshold of 0.01. The lower distance threshold for interaction between any two loci was 20kb and the upper distance threshold was 2mb; interactions spanning a distance outside this range were not considered for statistical significance.

### Loop Overlap and Aggregate Peak Analysis

For overlap between two sets of loops we used either no slack (comparison of normalization methods) or a slack of +/− 1 bin (alignment pipeline comparison). For aggregate peak analysis (APA), we used the APA function from the GENOVA (van der Weide et al., 2021) R package. APA score was computed as the value of the center pixel divided by the mean of pixels 15-30 kb downstream of the upstream loci and 15-30 kb upstream of the downstream loci (Phanstiel et al., 2015). APA ratio was computed as the ratio of central bin to the remaining matrix. For the alignment method comparison analysis, APA was performed for all loops and the top 10,000 significant FitHiChIP loops by q-value called using each alignment method (HiC-Pro, Juicer, or distiller-nf) with respect to the ICE-normalized contact matrix from each respective alignment tool. For the HiCCUPS normalization comparison analysis, APA was performed for the top significant HiCCUPS loops by the “donut FDR” for each normalization method (SCALE, VC, VC_SQRT) using the HiC-Pro-generated ICE-normalized contact matrix as background.

### Comprehensive Quality Control (QC) of HiChIP and ChIP-seq Samples

To evaluate sample quality and assign QC flags to all HiChIP (unmerged and merged biological replicates) and ChIP-seq samples, we selected and aggregated a comprehensive set of quality metrics from each processing step: read alignment, peak calling, and loop calling. For FH loops, HiChIP pre-processing was evaluated by number of reads, number of valid pairs, mean mapping percentage, and percentages of valid pairs, duplicate pairs, cis pairs, and cis long-range pairs, HiChIP peak calling was evaluated by number of peaks, and loop calling was evaluated by number of loops (**Supp. Table S8**). For FC loops, HiChIP read alignment, ChIP-seq peak calling, and loop calling as previously described (**Supp. Table S8)**. ChIP-seq pre-processing was evaluated by NRF (non-redundant fraction), PBC1 (PCR bottlenecking coefficient 1), PBC2 (PCR bottlenecking coefficient 2), NSC (normalized strand cross-correlation coefficient), and RSC (relative strand cross-correlation coefficient) (**Supp. Table S4**).

For each individual metric, QC scores were derived as follows. Based on the distribution of a given metric, along with field standards, we established three numerical intervals that are qualitatively considered “Poor”, “Warning”, and “Good” for that metric defined by thresholds *t*_1_ and *t*_2_ (**Supp. Table S9**). For each metric, we assigned samples a score between 0 and 10, with the 0 to 6 score interval representing “Poor”, 6 to 8 corresponding to “Warning”, and 8 to 10 to “Good”, by performing piecewise linear normalization of metric values. Samples were assigned a score of 10 if their metric value reached the maximum value (for bounded metrics) or was an upper outlier (exceeded the 75th *percentile* + 1.5 * *InterquartileRange*). Otherwise, the metric value *x_i_* for sample *i* was linearly re-scaled using a weight defined by 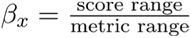, where:

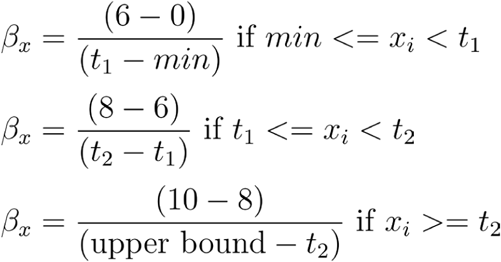

The metric score for sample *i* was then defined as *β_x_* * *x_i_*. For metrics for which higher values corresponded to poorer quality, such as duplicate pairs, this score was adjusted by calculating the complement 10 - *score_i_*. To aggregate individual metric scores from one processing step, such as read alignment, we calculated an average of the scores:

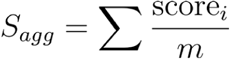

where *m* is the number of metrics used. Samples were assigned a QC flag for each processing stage based on *S_agg_* as follows:

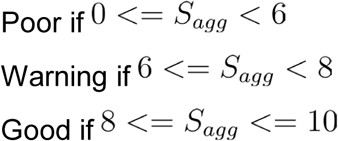

A final QC flag of “Poor”, “Warning”, or “Good” was derived for each configuration of loop calling (stringent 5kb, loose 5kb, stringent 10kb, loose 10kb, stringent 25kb, loose 25kb based on processing stage-specific QC flags (**Supp. Figure S4**). “Good” was only assigned in cases where all stage-specific QC flags were “Good”. All other possible cases are further described in **Supp. Table S9**. The final QC flag assigned to the merge filtered loops is equivalent to that of the non-merge filtered loops for the same sample.

QC flags are displayed within the main data table and on individual sample pages for visual inspection. Additionally, QC flags are automatically downloaded from the main data page (via the “CSV” button), even when the corresponding columns are not visible.

### Establishing Sets of High-Confidence HiChIP Samples

We established a high-confidence set of 54 unique human H3K27ac HiChIP samples from the pool of 156 human H3K27ac HiChIP datasets (merged or unmerged biological replicates) with either “Good” or “Warning” QC flags and over 10,000 stringent 5kb FC loops, henceforth called the High Confidence Regulatory Loops (HCRegLoops-All) sample set. We additionally established two subsets of the HCRegLoops-All sample set as follows: HCRegLoops-Immune contains 27 samples from immune-associated cell types and HCRegLoops-Non-Immune contains 27 samples from non-immune cell types (**Supp. Table S10**). Additionally, we defined a CTCF HiChIP sample set of 11 unique samples from the pool of 30 CTCF HiChIP datasets (merged or unmerged biological replicates) with either “Good” or “Warning” QC flags and over 2,000 stringent 5kb FC loops which we call the High Confidence Structural Loops (HCStructLoops) sample set (**Supp. Table S10**).

### Identifying Significant Chromatin Loops from High Resolution Hi-C Data

Loop calling was performed for 44 high resolution Hi-C samples gathered from the “High-Resolution Hi-C Datasets” Collection of the 4DN Data Portal (https://data.4dnucleome.org) using Mustache (v1.0.1) (Roayaei Ardakany et al., 2020). Processed .hic files were downloaded and Mustache was run with default parameters across chr1 through chr22 using raw, KR, and VC normalized contact matrices (-norm) to determine loops at 1kb, and 5kb resolutions (-r) (**Supp. Table S11**).

### Identifying SGLs in Immune-based Diseases

SGLs were identified utilizing fine-mapped GWAS SNPs from CAUSALdb for Type 1 Diabetes (T1D), Rheumatoid Arthritis (RA), Psoriasis (PS), and Atopic Dermatitis (AD) which included 7, 7, 3, and 1 individual studies, respectively (**Supp. Table S12**). The fine-mapped data was downloaded and lifted over from hg19 to hg38 using the MyVariantInfo Python package (version 1.0.0) (Xin et al., 2016). We downloaded the GENCODE v30 transcriptome reference, filtered transcripts for type equal to “gene” and located coordinates of the transcription start site (TSS) (Frankish et al., 2021). For genes on the plus strand, the TSS would be assigned as a 1bp region at the start site and, for the minus strand, the 1bp region at the end site. Lastly, we extracted all HiChIP samples whose organ was classified as “Immune-associated”. To integrate these datasets, loop anchors were intersected with fine-mapped GWAS SNPs and TSSs independently using bedtools *pairtobed*. Subsequently, loops were extracted as an SGL if at least one anchor contained a GWAS SNP and the opposing anchor contained a TSS.

### Calculating Significance for the Number of SGL Links

To assess the number of links we find through SGL mapping in comparison to linking all nearby genes, we built a null distribution. We downloaded all associations from the GWAS Catalog (1.0) (https://www.ebi.ac.uk/gwas/docs/file-downloads), filtered for variants with a p-value less that 5e-8 and linked each SNP to gene within 1Mb in either direction. A null distribution for the number of SNPs linked to genes, or vice versa, was calculated. We then used a left-sided Mann-Whitney test (scipy.stats.mannwhitneyu with alternative=’less’) to assign the probability that the number of linked SNPs or genes was less (i.e., more specific) for the SGL compared to the null distribution from GWAS Catalog.

### Identifying SGLs with Immune-associated eQTL Studies

SGLs were identified utilizing eQTLs from the eQTL Catalog which included CD4+ T cells, CD8+ T cells, B cells, natural killer (NK) cells and monocytes (**Supp. Table S13**). Similarly to GWAS-SGLs, we used GENCODE v30 to locate the TSS and extracted a subset of the GWAS-SGL HiChIP samples whose cell type matched the eQTL studies. To integrate these datasets, loops were intersected with pairs of SNP-gene pairs using bedtools *pairtopair*. Subsequently, loops were extracted as an SGL if at least one anchor contained an eQTL SNP and the opposing anchor contained a promoter.

### Building a Consensus Gene List for T1D

From the MalaCards database (https://www.malacards.org/), a query was made using the term “*type_1_diabetes_mellitus”* and the list of associated genes was downloaded. For eDGAR (https://edgar.biocomp.unibo.it/gene_disease_db/), “*diabetes mellitus, 1*” was queried under the Main Tables tab and all gene symbols were extracted. OpenTargets hosts a disease based search with gene association scores and a query was made to the MONDO ID for T1D (MONDO_0005147). The corresponding genes were downloaded and filtered for an association score > 0.5. For the GWAS Catalog, a query was made using the previous MONDO ID and all associated genes were extracted. Table 1 of Klak et al (Klak et al., 2020) summarizes genes that have been associated with T1D and gene symbols were extracted from this resource.

### Performing Motif Enrichment Analysis across Conserved Anchors

Motif enrichment analysis performed on conserved loop anchors from the HCRegLoops-All, HCRegLoops-Immune, HCRegLoops-Non-Immune, and HCStructLoops sample sets respectively. Briefly, for each sample set, we identified highly conserved loop anchors by aggregating the merge-filtered 5kb FC Loops across all samples in the set, extracting unique anchors, and selecting anchors involved in at least one loop call in at least approximately 80% of samples from the sample set (44/54 samples for HCRegLoops-All, 22/27 samples for HCRegLoops-Immune and HCRegLoops-Non-Immune, and 9/11 for HCStructLoops. Conversion from BED to FASTA was performed using the MEME *bed2fasta* utility (5.5.0). Repeats were masked using TRF (4.10.0) (Benson, 1999) with a match weight of 2, mismatch penalty of 7, indel penalty of 7, match probability of 80, indel probability of 10, minscore of 50, and maxperiod of 500 with the options -f -h -m specified. A biologically relevant background set of sequences was designed by first aggregating all non-conserved loop anchors, converting from BED to FASTA, and performing repeat masking. The non-conserved anchors were then downsampled to the number of conserved anchors using stratified sampling to match the GC content of the foreground conserved anchors. In short, non-conserved anchors were assigned to 0.5% quantiles based on GC content, and for each conserved sequence, GC content was computed and a non-conserved anchor from the corresponding quantile was sampled. We downloaded 727 known human motifs from the 2022 JASPAR CORE database (Castro-Mondragon et al., 2022). Motif enrichment analysis was directly applied to the conserved anchor sites using the downsampled non-conserved anchor sites as background using MEME Suite Simple Enrichment Analysis (SEA) (5.5.0) (Bailey et al., 2015) with a match e-value threshold of 10 (**Supp. Table S14**).

### Identifying Pairs of Motifs Overlapping Loop Anchors

Briefly, for each sample in HCRegLoops-All and HCStructLoops, unique merge-filtered 5kb FC loop anchors were extracted and these were intersected with sample-specific corresponding ChIP-seq peaks using *bedtools intersect* with no slack. We selected the peak with the highest signal value to represent that anchor for cases in which multiple H3K27ac peaks overlapped one loop anchor. The resulting peak sets were deduplicated since a single peak may overlap multiple loop anchors. To determine the genomic coordinates of motifs from the 2022 JASPAR CORE database (n = 727 motifs) in these representative peak sets, MEME Suite Find Individual Motif Occurrences (FIMO) (5.5.0) (Grant et al., 2011) was applied to the HCRegLoops-All sample set on a sample-by-sample basis using a match q-value threshold of 0.01 and default values for all other parameters. For each sample, we intersected motif coordinates with loop coordinates using *bedtools pairtobed* and annotated loop anchors with the motifs falling within the anchor.

### Performing Enrichment Analysis for Motif Pairs

Statistical analyses were performed via a bootstrapping method for HCRegLoops-All HiChIP samples (**Supp. Table S15, Supp. Figure S5**). To avoid shuffling with unmappable or repeat regions, we excluded loops where either anchor overlapped blacklisted regions from https://github.com/Boyle-Lab/Blacklist/blob/master/lists/hg38-blacklist.v2.bed.gz. We designed a block bootstrap approach with anchors as the unit analysis and anchors were randomly shuffled within their corresponding chromosomes. To then build a null distribution, we performed 100,000 simulations where we assigned a uniform probability of drawing a given anchor within each chromosome with replacement. P-values were determined for each sample by counting the total number of simulated pairs greater than or equal to observed pairs in the original sample. Due to the large number of pairs ranging from tens to hundreds of thousands for each sample, we focused our analysis on motif pairs within the top 50 most frequently enriched motifs for that sample. A multiple testing correction using Benjamini-Hochberg was applied to these filtered pairs to obtain adjusted p-values.

### Constructing Chromatin Interaction Networks using Loops

To construct a network from chromatin loops, each anchor was considered a node and each significant loop an edge. In addition, anchors were labeled as promoters by intersecting with TSS coordinates (slack of +/− 2.5kb) and allowing the promoter label to take priority over any other possible label. For H3K27ac HiChIP data, non-promoter nodes/anchors are labeled as enhancers when they overlap with ChIP-seq peaks (no slack). All other nodes that are not a promoter or an enhancer were designated as “other”. After obtaining the annotated anchors, we created a weighted undirected graph using the loops as edges and loop strength as edge weights calculated as the -log10(q-value) of FitHiChIP loop significance. To trim outliers, we set values larger than 20 to 20 and further scaled these values to between 0 and 1 for ease of visualization.

### Detecting and Prioritizing Communities

Community detection was applied to the networks created using FC loops at 5kb. Two levels of community detection were applied, the first detected communities within the overall network created for each chromosome (high-level) followed by a second round that detected subcommunities of the communities reported in the first round (low-level). High-level communities were located by running the Louvain algorithm using default parameters as implemented by the CDlib Python package (v0.2.6) (Rossetti et al., 2019). Starting with each high-level community, subcommunities were called using the same Louvain parameters. CRank (Zitnik et al., 2018) was then applied at both levels to obtain a score that aggregates several properties related to the connectivity of a community into a single score for ranking.

### Deriving 2D Models from Chromatin Conformation Data

To build 2D models we followed the framework laid out by the METALoci preprint (Mota-Gómez et al., 2022). We utilized 5kb resolution Hi-C data normalized with Vanilla Coverage and for each gene, we extracted the Hi-C matrix of +/−2Mb region surrounding that gene. For each such 4Mb region, we performed the following: 1) transformed normalized contact counts using log10, 2) identified contacts with low counts (log10 transformed value of <0.1 for HiChIP and <1 for Hi-C) and set them to zero, 3) derived a distance matrix by taking the reciprocal of the contact counts. Then, we extracted the upper triangular matrix and applied the Kamada-Kawaii layout to the graph represented by this matrix using networkx.kamada_kawai_layout(G). This process assigned a physical 2D location to each node and thus generated a 2D model (Kamada & Kawai, 1989). For visualization, all 2D models are visualized by overlaying the nodes with their contact count to the gene of interest for this 4Mb region.

### Analyzing 2D Models Using Spatial Autocorrelation Analyses

To analyze clustering of 1D epigenetic signals within a 2D model of chromatin we used spatial autocorrelation analyses via Global and Local Moran’s I as implemented in the ESDA (Exploratory Spatial Data Analysis) Python package (Rey & Anselin, 2007) and described in the METALoci methodology (Mota-Gómez et al., 2022). For HiChIP samples, the 1D signal was derived from corresponding ChIP-seq data. In the case of Hi-C, we processed the high resolution 4DN data with matching ChIP-seq signal data (**Supp. Table S16**). To focus this analysis, we extracted contact submatrices encompassing ±1Mb around each gene and started by obtaining Voronoi volumes using the Kamada-Kawai layout where each anchor became a node and, when interactions were present, edges were established using the interaction count as strength scores. 1D epigenetic signals were then overlaid onto the Voronoi polygons followed by calculation of spatial weights. The Global Moran Index was then applied to these spatial weights which statistically quantifies whether similar values are clustered or dispersed across space using esda.Moran(y, w). To identify specific clusters at a local level, we used the aforementioned spatial weights with the Local Moran Index—by applying esda.Moran_Local(). The resulting moran_loc.p_sim[i] provides information of the statistical significance (p-value) of each subregion; and moran_loc.q[i] shows the significance levels of each subregion: HH, HL, LH, and LL. For the significance levels, the first character symbolizes whether the node has a high (H) or low (L) signal value, and the latter character symbolizes whether its neighboring nodes have high or low signal values. To visualize the 2D model we start by plotting the Voronoi polygons overlaid with interaction between our gene of interest and all other regions (**Supp. Figure S6 top**) followed by two additional Voroni visualizations, one overlaid with ChIP-seq signals and the other with significance values from local Moran’s I, respectively (**Supp. Figure S6 middle**). Lastly, significance values for local Moran’s I are also visualized as a scatter plot to show the distribution of each anchor’s Local Moran Index score (**Supp. Figure S6 bottom**). Like the previous section, we provide visualizations of these results in the form of 2D models overlaid with interactions targeted to our gene of interest or overlaid with ChIP-seq signals and queried by gene names.

### Designing the Internal Database and Filesystem

The database is composed of two main parts, the first contains high sample-level and low-level loop information while the second contains SGL specific tables with auxiliary tables used to add additional annotations. For the first part, we atomized the data into the following tables: *hic_sample, celltype, hicpro, chipseq_merged, fithichip_cp, fithichip_cp_loop, hiccup, fithichip_hp* and *reference* which allowed us to capture important metadata and the uniqueness of loops using different loop callers. The second part includes the *gwas_study, gwas_snp, snp, gene*, *fcp_fm_sgl*, eqtl_study, eqtl, and fcp_eqtl_sgl tables which allowed us to capture important relationships for SGLs and to facilitate their query. The full database schema can be found in **Supp. Figure S7.**

### Designing and Implementing the Web Interface

The Loop Catalog was built using Django (3.2) (https://www.djangoproject.com/) as the backend framework with all data stored using the Postgresql (9) (https://www.postgresql.org/) database management system. To style the frontend interface, we used Bootstrap (5.2) (https://getbootstrap.com/docs/5.2/). For implementing advanced tables and charts DataTables (1.12.1) (https://datatables.net), Charts.js (4.0) (https://www.chartjs.org/), and D3 (4.13.0) (https://d3js.org/) were used. To visualize genetic and epigenetic data the WashU Epigenome Browser (53.6.3) (https://epigenomegateway.wustl.edu/) was used which maintains an easily accessible web embedding. Lastly, CytoscapeJS (3.26.0) (https://js.cytoscape.org/) was used to visualize enhancer-promoter networks.

## UTILITY AND DISCUSSION

### Curating a set of publicly available HiChIP samples for human and mouse cells

In total, we collected 763 distinct human and 281 distinct mouse samples from 152 studies (1334 total with replicate-merged samples included) that cover a diverse set of cell types and cell lines. For primary human samples there is a concentration of immune cell types that include monocytes, natural killer cells, T cell and B cell subsets, among others (237 samples designated as immune associated; 186 and 51 for human and mouse, respectively). As expected, cancer cell lines are well represented (e.g., HCC15, NCI-H1105, MCF7) together with other cell lines from normal tissue including heart-derived samples (e.g., aortic valve interstitial, coronary artery smooth muscle, and aortic smooth muscle cells) as listed in **Supp. Table S1**.

Regarding the target protein in the HiChIP experiment, active regulatory element-associated histone mark H3K27ac was the most highly represented, accounting for ~60% of human samples and ~54% of mouse samples. Other frequently represented ChIP targets include CTCF for human datasets and H3K4me3 for mouse datasets. Cohesin subunits such as SMC1A was also a frequent protein of choice in both human and mouse samples (**Figure 1C**). It is evident that, while the majority of studies target functional/regulatory interactions via H3K27ac or H3K4me3 pulldown, structural interactions via CTCF and cohesin pull-down are also well-represented among human and mouse HiChIP studies. As will be further discussed below, we processed our HiChIP and auxiliary data such as ChIP-seq, when available, using different methods for peak calling (**Supp. Figure 8A-B**) and loop calling including loops at three different resolutions to provide users options for different levels of depth for downstream analysis (**Supp. Figure 8C-D**). Overall, this dataset provides a comprehensive coverage of HiChIP samples that investigate structural as well as regulatory loops profiled across hundreds of samples.

### Uniform processing of HiChIP samples and quality controls

#### Alignment Pipelines

We first tested multiple different pipelines that are commonly used for HiChIP data processing including HiC-Pro, Juicer, and distiller-nf on a subset of 45 human samples from diverse cell types and protein pulldowns. Juicer and distiller-nf reported a comparable number of valid pairs, cis interaction pairs, cis short-range and long-range interaction pairs, and trans interaction pairs to HiC-Pro (**Supp. Figure S9A-B**). Next we assessed the sets of loops derived from these three different pipelines. There was a good, although not perfect, overlap (~81%) among loops reported by these three pipelines (**Supp. Figure S9C**). HiC-Pro-reported valid pairs demonstrated stronger enrichment in aggregate peak analysis (APA) compared to those derived from Juicer or distiller-nf pairs (**Supp. Figure S9D**). We, therefore, proceeded with HiC-Pro and aligned our full collection of HiChIP data to human (hg38) or to mouse (mm10) genomes using appropriate configuration (e.g., restriction enzyme) for each sample.

#### Peak calling

Next, we assessed possible ways to call peaks of the 1D HiChIP signal which becomes essential in the absence of matched ChIP-seq data (MACS2 was used for ChIP-seq peak calling). We previously developed a module in FitHiChIP (also utilizes MACS2) for peak calling from HiChIP data (Bhattacharyya et al., 2019), and another tool named HiChIP-Peaks was also developed for the same purpose (Shi et al., 2020). When we compared these two approaches, peaks called by HiChIP-Peaks spanned uncharacteristically large genomic regions with median per-sample average peak size of 2.26kb (for reference, ~2.3 billion peaks from ChIP-Atlas had a median size of 304bp). Furthermore, peaks from HiChIP-Peaks achieved approximately half the percent recall of ChIP-seq peaks compared to that achieved by FitHiChIP peaks for the same total peak span (**Supp. Figure S2A-B**). HiChIP-Peaks is additionally not compatible with the MNase HiChIP samples represented in the Loop Catalog. Thus, we proceeded with FitHiChIP-based peak calls for downstream analyses.

#### Loop calling

For the identification of loops from HiChIP data, there again are a number of computational methods available, including those originally designed for Hi-C (e.g., HiCCUPS) and others specifically for HiChIP (e.g., FitHiChIP, hichipper, MAPS) (Bhattacharyya et al., 2019; Juric et al., 2019; Lareau & Aryee, 2018). Us and others have previously compared these tools in detail showing specificity and sensitivity trade-offs (Bhattacharyya et al., 2019; Juric et al., 2019; Lareau & Aryee, 2018) that are important to consider for different downstream analysis tasks in hand. In this work, we employed HiCCUPS to represent a highly specific loop caller and FitHiChIP to represent a method with higher sensitivity. In total, we utilized three different approaches for the identification of significant loops: **i)** FitHiChIP with ChIP-seq peaks (FC loops), **ii)** FitHiChIP with peaks inferred from HiChIP by the FitHiChIP *PeakInferHiChIP.sh* utility (FH loops), and **iii)** HiCCUPS (**Supp. Figure S1**).

#### Contact map normalization

Another important consideration for HiChIP loop calling is the contact map normalization. For instance, HiCCUPS is compatible with multiple normalization approaches including KR (Knight and Ruiz), VC (Vanilla Coverage), VC_SQRT, and SCALE. Although commonly used for Hi-C datasets, KR normalization frequently did not converge for one or more chromosomes at all resolutions (of the 45 HiChIP samples, only 12 (26.6%), 11 (24.4%) and 12 (26.6%) samples had all autosomes completing KR normalization step for 5kb, 10kb and 25kb resolutions, respectively) (**Supp. Table S17**). VC, VC_SQRT, and SCALE converged for all chromosomes at all resolutions for these 45 samples (**Supp. Table S17**). When we assessed multiple normalizations coupled with HiCCUPS, VC_SQRT normalization reported a surprisingly large number of loops across sample (median of 26,507) that was quite different than that of VC (median of 5380) and SCALE (median of 5052) (**Supp. Figure S10A**). When we assessed overlap of 5kb HiCCUPS loop calls for two samples (H9 embryonic stem cells and CD34+ cells from cord blood), only ~19% were commonly reported by VC, SCALE and VC_SQRT normalizations largely due to 5-fold higher loop calls by VC_SQRT (**Supp. Figure S10B**). Evaluating the loop calls further using APA showed that VC led to highest enrichment among the three normalizations for both samples (**Supp. Figure S10C**). Thus, we ran HiCCUPS on all HiChIP samples using VC normalized contact matrices.

#### Hi-C loops

In addition, we performed loop calling using Mustache (Roayaei Ardakany et al., 2020) for 44 high-resolution Hi-C samples available from the 4DN data portal at 1kb and 5kb resolutions using raw, VC, and KR normalizations. When focusing on the set of loops called utilizing KR normalized counts, we obtained on average ~48k loops for both 1kb and 5kb resolution across all Hi-C samples (**Supp. Table S11**).

#### Loop visualization

To visualize Hi-C and HiChIP loops, we developed the Loop Catalog webpage, which includes an interactive table that allows selection of multiple samples, loop calling settings, resolutions and corresponding peaks (**Figure 2**). These selections can then be visualized in an embedding of the WashU Epigenome Browser within the Loop Catalog website or downloaded as a track file or WashU hub file for external use. Alternatively, if users would like to stay within the Loop Catalog ecosystem, session files can be downloaded and uploaded to save a snapshot of their work for later use (**Figure 2B**).

**Figure 2.**
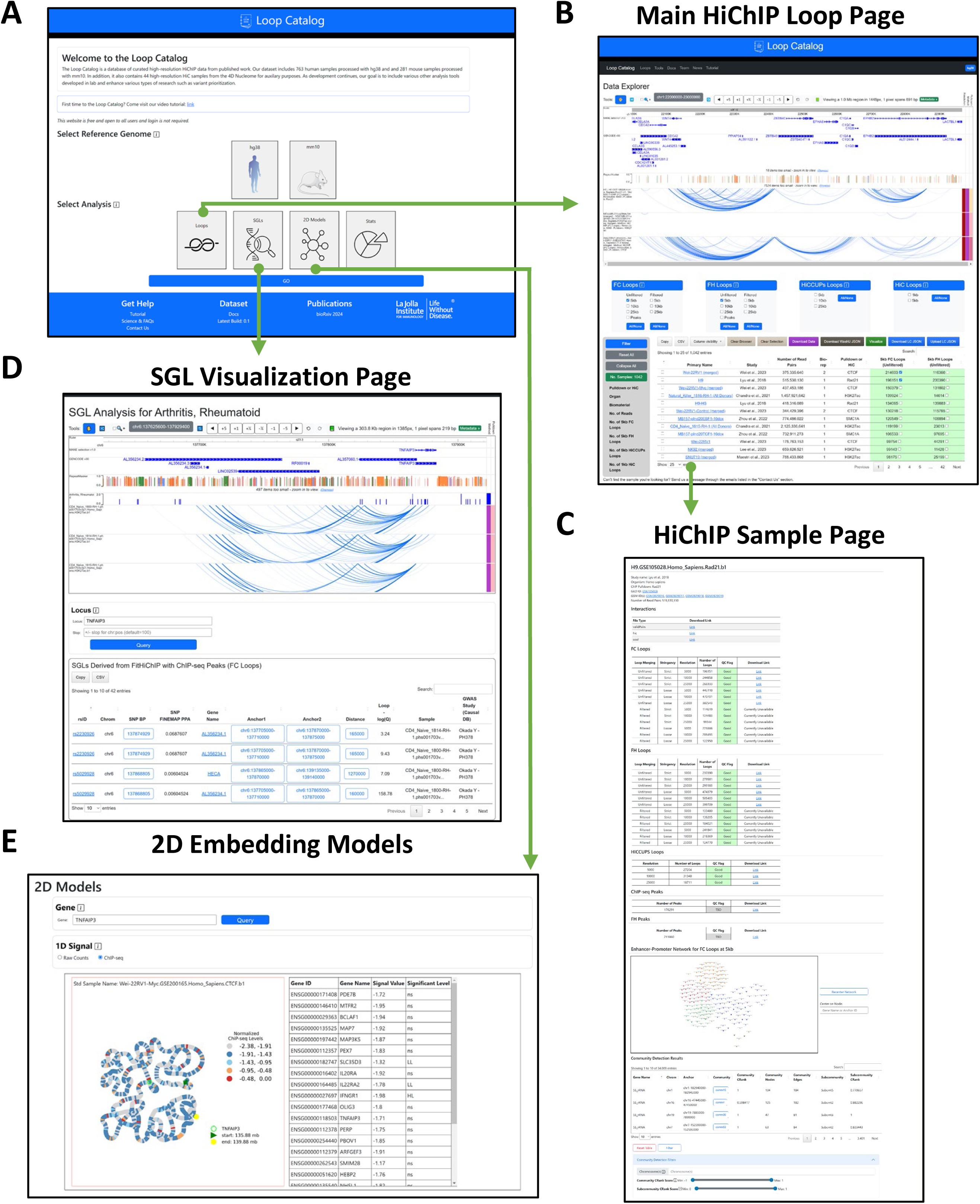
Layout of the Loop Catalog portal. **A)** Screenshot of the main entry page to the Loop Catalog. **B)** Main data page which includes an embedding of the WashU Epigenome Browser followed by a table of HiChIP samples with various metadata fields. **C)** Screenshot of a HiChIP sample page with download links, summary of loop call statistics and an enhancer-promoter network visualization with an accompanying table listing detected community and subcommunity’s of this network (enhancers - circles, promoters - squares and other regions - triangles). **D)** Screenshot of the GWAS-SGL page with the locus of interest centered and a table of SGLs with navigation buttons. **E)** Screenshot of the 2D embedding models page which includes a visualization for each sample for the queried gene locus, a table of spatial autocorrelation analysis, and buttons to swap between 1D overlap of ChIP-seq or interaction (raw) signals.

#### Data summary

The Loop Catalog was initialized with loops called for 1044 unmerged HiChIP samples and 284 merged samples that were created after combining biological replicates from the same study that pass a correlation threshold for pairwise similarity of individual replicates (**Supp. Figure S3**). An additional 6 immune cell-based *mega-merged* samples were created by merging across all donors and all biological replicates using data from two previous publications (Chandra et al., 2021; Schmiedel et al., 2021). Overall, the Loop Catalog provides access to loops for 1334 HiChIP samples with FH loop calls for all samples and, of these, 386 samples had matching ChIP-seq data that enable loop calling using FC. In addition, we called genome-wide HiCCUPS loops for 442 samples for which the number of 10kb loop calls from chr1 only passed our thresholds (200 for human, 100 for mouse). Loop Catalog encompasses HiChIP loops from a diversity of protein pulldowns that exhibit different characteristics such as high (e.g., H3K27ac) or low (e.g., H3K9me3) number of loops, in general, for different histone modifications (**Supp. Figure S11**). In addition to the number of loops, the characteristic loop size (i.e., genomic distance between anchors) also varies depending on the protein pulldown. This is the case with loops from CTCF or cohesin complex subunits (median size of 190kb and 175kb) compared to H3K27ac loops (median size of 115kb) for human samples and with similar results in mouse (**Supp. Figure S12**).

#### Comparison to other databases

When comparing the Loop Catalog to other databases, the Loop Catalog stands out with the highest number of HiChIP samples, totaling 1044 distinct and 1334 overall with merged samples. ChromLoops (Zhou et al., 2023) provides loop calls for 816 samples (772 HiChIP); however, this total spans 13 species. HiChIPdb (Zeng et al., 2023) and CohesinDB (J. Wang & Nakato, 2023) offer significantly fewer HiChIP samples, with only 200 and 42, respectively (**Table 1**). It is noteworthy that different loop calling methods were preferred by different databases. With this consideration, HiChIPdb is most similar to the Loop Catalog, employing FitHiChIP for loop calling (**Table 1**). However, unlike the Loop Catalog, HiChIPdb derives peaks solely from HiChIP data, which tend to have a low recall rate with respect to ChIP-seq peaks (**Supp. Figure S2**). Conversely, ChromLoops opted to use ChIA-PET Tool (V3) (G. Li et al., 2019), a tool specifically designed for ChIA-PET (chromatin interaction analysis with paired-end tag) experiments and may not be ideal for HiChIP. Cohesin-DB employed HiCCUPS for loop calling, a feature of our database also, which comes with previously discussed limitations about low sensitivity and low applicability (i.e., a large fraction of samples with zero or very small number of loop calls) (Durand et al., 2016) (**Table 1**). Another important advantage of the Loop Catalog is the rigorous quality control (QC) performed for all stages of processing, including read alignment, peak calling (ChIP-seq or HiChIP-derived), and loop calling, based on a comprehensive set of metrics (**Supp. Table S9**). We provide final QC flags for each set of loop calls to inform users of the inferred quality of the data they intend to visualize or download for further analysis (**Supp. Figure S4, Supp. Table S8**). Finally, Loop Catalog provides utilities including some previously offered by other databases such the ability to visualize loop calls onsite and integrate with GWAS variants and other utilities that are unique such as: 1) the integration of ChIP-seq peak calls with HiChIP data to derive FC loops, 2) GWAS-SGL and eQTL-SGL analyses with navigation across the different elements, 3) demonstration of database utility through motif enrichment analysis in conserved loop anchors across diverse cell types, 4) development of motif pair analysis for pairs of genomic regions, 5) availability of chromatin conformation networks with labeling of enhancers and promoters, and 6) generation of 2D embeddings models of chromatin conformation.

### Visualization and exploration of loop calls through a web interface

The Loop Catalog is underpinned by a comprehensive database that incorporates processed HiChIP data from GEO and dbGaP (**Figure 1, Supp. Table S7**), high-resolution Hi-C data from the 4DN data portal (**Supp. Table S11**), along with fine-mapped GWAS data for a number of immune-associated diseases (**Supp. Table S12**), eQTLs for major immune cell types (**Supp. Table S13**), and 2D models of chromatin conformation (**Supp. Figure S6**). When first accessing the platform, users are presented with an entry page featuring a selection for reference genome and analysis type including loops, SNP-Gene linking (SGLs), and statistics (**Figure 2A**). Subsequently, once an analysis has been selected, a secondary page will render the corresponding analysis with a navigation menu to switch between other analyses and website-related information. The data page offers immediate and extensive visualization of HiChIP and Hi-C samples with their associated loop calls, spanning various methods and resolutions, as illustrated in **Figure 2B**. For specific sample information, each sample is linked to a dedicated page displaying metadata, loop data, regulatory network analyses, and a motif scanning analysis (**Figure 2C**). In addition to visualization through Epigenome Browser, loops can also be visualized as networks of connected regulatory elements including enhancers and promoters. Topological structures such as communities and subcommunities in these networks alongside measurements of their connectivity properties (CRank) can be explored further through an interactive user interface. (**Figure 2C bottom**). Furthermore, the SGL page grants access to the integration of immune-based HiChIP samples and fine-mapped GWAS SNPs and eQTLs (**Figure 2D**). We also utilized METALoci (Mota-Gómez et al., 2022) to generate 2D embedding models, offering an alternative on chromatin conformation visualization. This approach simultaneously analyzes the spatial autocorrelation between 1D signals (e.g., ChIP-seq) within the model and is available as a separate module (**Figure 2E**). Lastly, the statistics page provides a higher-level overview of all HiChIP data stored for a given reference genome.

### Expanding the use of HiChIP data for annotating GWAS SNPs and eQTLs using SNP-to-Gene Loops

Leveraging the Loop Catalog, we further analyzed loops for immune-related HiChIP samples and how they overlap genetic variants and genes of interest. We identified 79 samples with nonzero FC loops at 5kb (63 unmerged, 10 from those merged at the biological level, and 6 all donors merged). We then used these loop calls together with fine-mapped GWAS SNPs from CAUSALdb to find potential target genes for each SNP that we term a SNP-gene pair with a loop (SGL) (**Figure 3A**). Briefly, across T1D, RA, PS and AD, there are 7729, 1121, 590, and 674 unique fine-mapped GWAS SNPs, respectively (**Figure 3B**). After overlapping these SNPs with our loop anchors and genes connected through such loops, we found 74 samples with at least one SGL and a total of 182,306 SGL instances across 18 studies covering the four diseases. When removing duplicate SGLs we found 28,162 distinct SGLs which included 4241 SNPs, 2354 genes and 7269 loops (**Figure 3B, Supp. Table S12**). To store these results into the database and to allow querying, we constructed SNP, GWAS, gene and loop-level tables with genomic coordinates. To then browse these results, an SGL entry page allows users to first select their target disease, locus and samples from which the loops will be derived (**Figure 2D**). Subsequently, the Loop Catalog returns an SGL analysis page that includes an embedded WashU Epigenome Browser element loaded with a track for fine-mapped SNPs and loops tracks for each sample. Below the browser users will find an interactive table that lists all mapped SGLs for their selection. This table also allows navigating between loops and within loops including the left or right anchor and SNP positions (**Figure 2D**).

**Figure 3.**
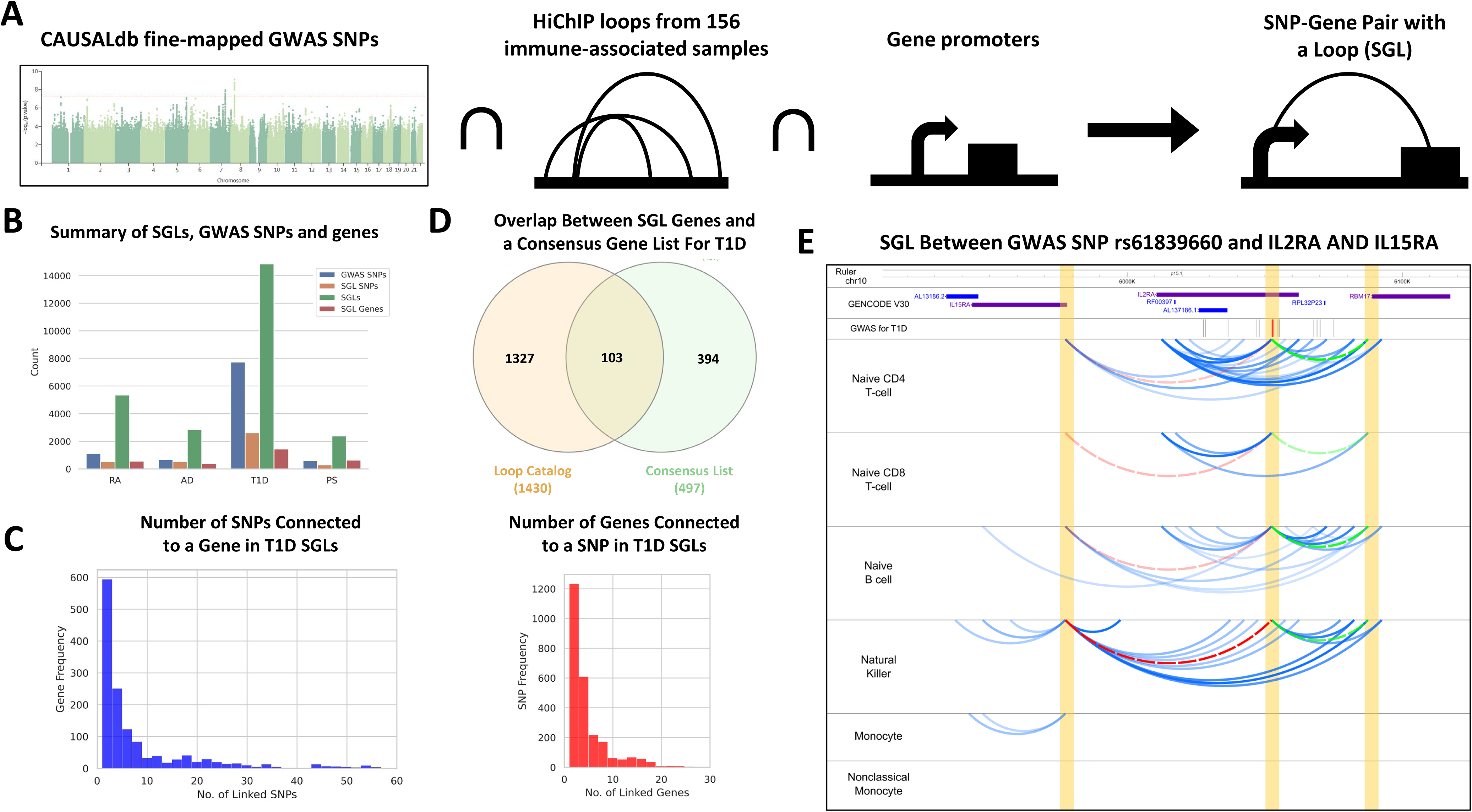
SGL analysis overview and results. **A)** Schema of the SGL analysis using fine-mapped SNPs from CAUSALdb, Loop Catalog immune-related samples, and TSS coordinates. **B)** Summary of results across all 4 diseases including the total number of GWAS hits (blue), SNPs found in a SGL (orange), genes found in SGL (green) and total SGLs (red). **C)** Distribution of SNP counts with respect to gene (left) and the distribution of gene counts with respect to SNP (right) for T1D. **D)** Evaluating the number of SGL genes which belong to a consensus list of T1D genes (green) and unique (orange). **E)** Example of an SGL between rs61839660 (red) and the genes IL15RA (red arc) and RBM17 (blue arc). Six tracks with arcs represent H3K27ac HiChIP loops for naïve CD4+ T cell, naïve CD8+ T cell, naïve B cell, Natural Killer, monocytes, nonclassical monocytes derived from the Schmiedel et al 2018 samples that were merged across all donors.

Similar to GWAS variants, we expanded our annotation of genomic elements using expression quantitative trait loci (eQTL) associations between SNPs and genes. We started by downloading uniformly processed eQTL studies from the eQTL Catalogue (Kerimov et al., 2021) and focused on cell types from Schmiedel et al. 2018 dataset that include eQTLs for naive CD4+ T cells (n=64386), naive CD8+ T cells (n=67793), naive B cells (n=60629), NK cells (n=48221), monocytes (n=66024) and nonclassical monocytes (n=58069). For these cell types, we also have the HiChIP data derived from a subset of the same donors (Chandra et al., 2021; Schmiedel et al., 2018, 2021). For these cell types, we then located 11604 unique eQTL-SGLs that cover 11053 SNPs and 1128 genes (**Supp. Figure S13**, **Supp. Table S13**). These results are made available through a similar web interface as SGLs derived from GWAS (**Supp. Figure S14**). It is possible to extend our eQTL-based SGLs analysis to the remainder of the eQTL Catalogue, however, matching cell types from eQTL studies to those from HiChIP studies is not a trivial task for most of the cases.

### Utilizing T1D SGLs for SNP and Gene Prioritization

To better understand the utility of using SGLs for linking GWAS SNPs and genes, we focused our attention on analyzing SGLs in T1D. When compared to all other diseases, T1D has the highest number of unique SGLs (n=16,534) mainly due to the high number of fine-mapped GWAS SNPs as our starting point (**Figure 3B**). In post-GWAS analyses, it is important to distinguish putative causal SNPs among those that are in linkage disequilibrium (high LD). In cases where the phenotypic effect of the SNP is mainly through regulation of a distal gene, SGLs can corroborate important information to accurately annotate SNP function and to prioritize GWAS genes and SNPs while utilizing information on their 3D proximity. Investigating this for T1D, we observed that at least half of the CAUSALdb SNPs participate in an SGL, these SNPs are often in contact with multiple genes (**Figure 3B**).

We further explored the multiplicity of SNP and gene links within T1D and found that 26% of SGL genes are linked to a single SNP, and the median number of SNP links per gene is 3. On the other hand, 27% of SGL SNPs are linked to a single gene and the median number of gene links per SNP is 2 (**Figure 3C**). To better understand this prioritization potential of SGLs we contrasted our methodology with a simple methodology of linking a GWAS SNP to all protein coding genes within +/− 100kb (nearby genes). This led to a median number of 3 genes per SNP, which was not far from 2 genes per SNP with SGLs. However, the median number of SNPs per gene was 7 for the “nearby genes” approach in comparison to 3 from SGLs leading to a statistically significant reduction (p-value <2.6e-61; MannWhitney U Test) suggesting better prioritization of relevant SNPs for each gene.

To understand if SGL genes overlap genes with known T1D associations, we built a consensus gene list using MalaCards (Rappaport et al., 2013), eDGAR (Babbi et al., 2017), OpenTargets (Ochoa et al., 2023), and GWAS Catalog (Sollis et al., 2023) and a T1D review paper (Klak et al., 2020). The union of the gene lists across these five resources had 497 genes in total, of which 106 overlapped with our 1532 SGL genes identified for T1D. The 1426 genes uniquely found by our SGL approach, although likely to involve false positives, are potential targets for future investigations (**Figure 3D**). One of these genes was *IL15RA*, a cytokine receptor that binds the pro-inflammatory cytokine IL-15 with high affinity and through *cis* and *trans* presentation of IL-15 impacts cellular functions of CD8+ T cells as well as Natural Killer (NK) cells (Waldmann, 2015). Despite *IL15RA* not being within the consensus T1D gene list, the IL15RA/IL-15 axis has been associated with T1D but whether this axis has a pathogenic or protective role has not been clear (Bobbala et al., 2017; J. Chen et al., 2013; Lu et al., 2020). Through our SGL analysis of H3K27ac HiChIP loop calls from major immune cells, we identified looping between the *IL15RA* promoter and rs61839660, a SNP 75kb away that is highly associated with T1D and has been further prioritized by fine-mapping in three out of four T1D-GWAS studies with a posterior probability greater than 0.70. The corresponding loops are found for T cells, Naive B cells and NK cells but not for monocytes suggesting an important role for this SGL within the adaptive immune system. The more likely scenario is that rs61839660’s T1D association is mediated through *IL2RA* given that this SNP falls within a constituent intron. However, specific loops connecting rs61839660 to *IL15RA* (P-value < 10^-9^) as well as to *RBM17* (P-value < 10^-11^) suggest the possibility of a pleiotropic effect for this SNP (**Figure 3E**, **Supp. Table S12**). In addition, *RBM17*, an RNA-binding protein, has been previously shown to affect other autoimmune diseases such as rheumatoid arthritis (Ibáñez-Costa et al., 2022). As exemplified here, SGL analysis with the Loop Catalog may provide further evidence and/or mechanisms of action for a genetic variant and its target gene. In addition, it may help to find targets for GWAS variants whose target gene remains elusive.

### Identifying Significant Sequence Motifs at Conserved Regulatory Loop Anchors

In order to examine binding patterns of TFs in regulatory loops, we performed 1D motif enrichment analysis on highly conserved regulatory loop anchors from three high-confidence (HC) sample sets of H3K27ac HiChIP samples, H3K27ac being a histone mark of active transcription (**Figure 4A**). The HCRegLoops-All sample set contains the 54 H3K27ac HiChIP samples with over 10,000 stringent 5kb FC loops. These samples encompass diverse cell types including immune cells, heart cells, and various cancer cell lines. The HCRegLoops-Immune sample set contains the subset of 27 samples from the HCRegLoops-All set which are from immune-associated cell types and the remaining 27 samples in HCRegLoops-Non-Immune contain a diverse set of cell types including multiple cancer cell lines. For the aggregate set of unique loops from each sample set respectively, we annotated a loop anchor as conserved if it was involved in at least one loop in 80% or more of the samples from the sample set. We identified 1160, 2410, and 1006 anchors fitting this criterion for HCRegLoops-All, HCRegLoops-Immune, and HCRegLoops-Non-Immune, respectively. Comparing the HCRegLoops-Immune and HCRegLoops-Non-Immune sample sets, while there were more unique loop anchors exclusive to HCRegLoops-Non-Immune, HCRegLoops-Immune had more exclusive conserved anchors, which is consistent with the higher degree of cell type similarity in HCRegLoops-Immune (**Figure 4B**). Using the Simple Enrichment Analysis tool SEA from the MEME Suite to perform motif enrichment analysis with 727 known human motifs from the 2022 JASPAR CORE database, we identified 204, 225, and 190 significantly enriched motifs (e-value < 0.01) for the HCRegLoops-All, HCRegLoops-Immune, and HCRegLoops-Non-Immune sample sets, respectively (**Supp. Table S14**). As a means of method validation, when the analysis was performed on the HCStructLoops sample set (CTCF HiChIPs), 5 enriched motifs were reported with an e-value < 0.01, CTCF being the top ranked motif with an enrichment ratio of 3.56 (**Figure 4C**). There was notable overlap between reported motifs discovered in conserved anchors from HCRegLoops-Immune and HCRegLoops-Non-Immune (**Figure 4B**) and in the top 15 motifs from all H3K27ac sample sets (**Figure 4C**). We believe that the aggregation of samples across diverse cell types may cause this loss of cell type-specificity in the reported motifs. Notably, among the H3K27ac sample sets, NRF1 was among the top 3 enriched motifs across all sample sets with over 2-fold enrichment (**Figure 4C**). Host cell factor C1 (HCF-1) has been shown to be involved in the formation of short-range loops between cis-regulatory elements (S. Liu et al., 2022) and is known to be a mutual binding partner of NRF1 (Han et al., 2017; Sekine et al., 2018), which may indicate that NRF1 has a role in gene regulation through chromatin loop formation in a cell type-independent manner. Our motif enrichment analysis also identified members of multiple zinc finger protein families among the top ranked motifs, such as EGR1, EGR2, EGR3, PATZ1, KLF15, SP1, and SP4, which are recognized as universal stripe factors, or proteins that bind to GC-rich sequences, increase DNA accessibility to binding partners, and lengthen residence time of colocalized factors (Zhao et al., 2022). PATZ1 was additionally recently shown to have novel chromatin insulating activities, similar to CTCF (Ortabozkoyun et al., 2022, 2023). However, recent studies highlight the importance of caution and the requirement for functional validation by showing that, for ZNF143, another zinc finger protein with a presumed role in looping (which has not come up as significant in our analysis), such an association was a result of antibody cross-reactivity (Magnitov et al., 2024; Narducci & Hansen, 2024). We have to note that since our analysis focuses on loop anchors conserved across many samples, we do not utilize cell-specific information (e.g., ATAC-seq or ChIP-seq peaks) to narrow down specific regions within an anchor (5kb), a common practice that is utilized for motif enrichment analysis to pinpoint cell-type specific transcription factors. Therefore, it was expected to observe enrichment of motifs that bind non-cell-type-specific TFs (e.g., ZNF family members). While future work is needed to validate the TFs with enriched motifs identified here, our analysis of loop anchors conserved across a diverse set of samples is useful for developing hypotheses about generic regulators or correlates of looping.

**Figure 4.**
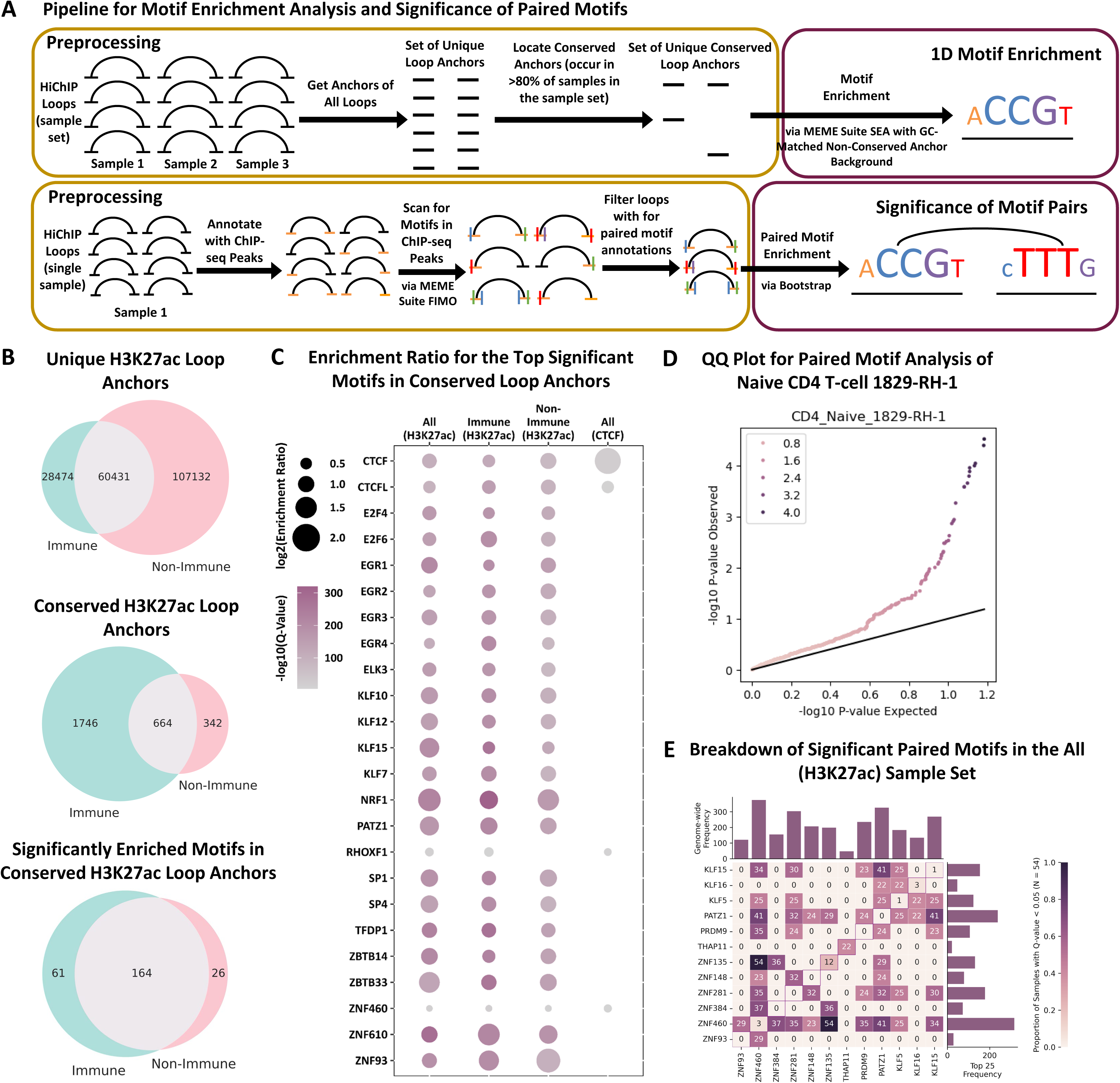
Motif and paired-motif analysis of loop anchors. **A)** Schematic of the 1D and 2D (paired) motif analysis. For 1D motif enrichment analysis, HiChIP loops are aggregated across all samples in the sample set, conserved anchors are identified, and motif enrichment analysis is performed directly on the loop anchors. Paired motif analysis is performed on a per-sample basis. HiChIP loops from a single sample are overlapped with ChIP-seq peaks and paired motif analysis for loops is applied after motif scanning in ChIP-seq peaks. **B)** For the 1D motif enrichment analysis, Venn diagrams show the overlap between immune (n = 27) and non-immune (n = 27) H3K27ac HiChIP sample sets for unique loop anchors, conserved loop anchors, and significantly enriched motifs in conserved loop anchors reported by MEME Suite SEA (e-value < 0.01). **C)** Bubble plot for the union of the top 15 motifs from each H3K27ac sample set (HCRegLoops-All, HCRegLoops-Immune, and HCRegLoops-Non-Immune) and top 6 motifs from the CTCF sample set (HCStructLoops) (24 total motifs). The q-value is represented on a range from 0 (gray) to 300 (magenta) and the log2(enrichment ratio) is represented by a circle radius from 0.5 to 2. **D)** QQ plot testing p-values for Naive CD4+ T cell 1829-RH-1 for the paired motif analysis bootstrap method. **E)** Heatmap of significant motif pairs (center) where rows and columns represent motifs on opposite anchors and each cell represents the proportion of samples where the given motif pair is significant. The distributions of a given motif across the whole genome and within the top 25 motif pairs are represented on the top and right, respectively.

### Identifying Significant Motif-Pairs Across Loop Anchors

Transcription factors often work together, through co-binding, dimerization, or multimerization, to promote gene expression (Bell et al., 2024; M. Li et al., 2023; Lupo et al., 2023; S. Rao et al., 2021; Rauluseviciute et al., 2024). With this motivation, we expanded our motif analysis at conserved anchors to a search for significantly overrepresented motif pairs connected by loops. Since the brute-force approach of scanning all possible motifs across the genome for enrichment would have been infeasible, we performed bootstrap analysis to calculate an empirical p-value. We used the frequency of a given motif pair across the entire set of loops as our statistic and tested whether the frequency of this motif pair appeared greater than expected by chance for a given loop set. Bootstrapping was used to build a simulated null distribution by intra-chromosomal shuffling of loop anchors and thus motif pairs (see methods “Performing Enrichment Analysis for Motif Pairs”). A p-value was then calculated as the fraction of simulations greater than our observed frequency (**Figure 4A** (bottom); **Supp. Figure S5**). We performed this analysis on the HCRegLoops-All sample set (**Supp. Table S15**) and, to ensure our bootstrapping approach did not lead to inflated p-values, we investigated the Q-Q plots of our samples. We observed that most p-values lie along the diagonal for non-significant p-values suggesting no inflation (**Figure 4D**). Summarizing this analysis across all of our samples, we found signals for ZNF460 paired with motifs such as ZNF135 and PATZ1 across 100% and 75.9% of samples, respectively (**Figure 4E**). Pairs of PATZ1 and KLF15, both of which are universal stripe factors (Zhao et al., 2022), were also overrepresented across the majority of samples. Overall, this is an interesting finding which suggests an association for zinc finger proteins other than CTCF with chromatin loops and their anchor regions.

### Inferring 2D representations of chromatin structure from HiChIP data

Chromatin interactions are often depicted using 2D matrices, however, the matrix structure can abstract away the true physical and visual representation of a loop, TAD, or other chromatin conformation (T. Liu & Wang, 2018; MacKay et al., 2017). This limitation can be overcome by using 3D models of chromatin conformation, however, these require specialized methods to infer and interactive tools to visualize making them difficult to use for exploratory analysis or hypothesis generation. Another alternative, 2D models can compress the data into two dimensions while visually restoring some of the physical proximity and allowing easy overlay of other epigenetic signals (Mota-Gómez et al., 2022). To do just this, we applied METALoci to generate 2D models for HCRegLoop samples and 5 major cell lines with high resolution Hi-C. In addition to 2D modeling, METAloci analyzes spatial autocorrelation of overlaid 1D epigenetic signals. We applied METAloci analysis to HiChIP samples utilizing corresponding ChIP-seq data as the 1D signal and, similarly, utilized our Hi-C samples together with matched H3K27ac ChIP-seq signals for that sample (**Supp. Table S16**). More specifically, for each gene, we extracted submatrices encompassing ±1Mb around each gene, applied METALoci to generate 2D embeddings and generated the static images that represent these models alongside overlaid data. On our webserver, these sets of images can be browsed for a specific Hi-C or HiChIP sample using a query gene name adding a useful feature to the Loop Catalog (**Figure 2E**). As an example we investigated the genes *MCM3*, *CEP85L*, *FAM135A*, and *ESR1* within the mega-merged naive CD4+ T cell sample. The first two genes, *MCM3* and *CEP85L*, have been shown to be highly expressed in CD4+ T cells from blood (Schmiedel et al., 2018) and our corresponding 2D models revealed important enhancer activity that is spatially near to these genes of interest (**Supp. Figure S6A-B)**. In contrast, *FAM135A* and *ESR1* have low expression in CD4+ T cells (Schmiedel et al., 2018) and the 2D models for these genes showed markedly low levels of enhancer activity that localized close to these genes (**Supp. Figure S6C-D).**

## CONCLUSION

The HiChIP assay has empowered the field of chromatin structure by providing a relatively inexpensive, high-resolution and targeted approach to mapping chromatin interactions. It is no surprise that the number of HiChIP studies published annually keeps growing each year and will continue to do so until superior methods are developed. The Loop Catalog is a public hub that centralizes 1334 HiChIP samples from these studies and, through the use of user-facing tools, lowers the bar for exploring 3D genome organization datasets, which would require substantial bioinformatics skills otherwise. Loop Catalog offers the largest collection of HiChIP data while providing an extensive set of quality control criteria on the HiChIP data itself, which can be utilized by the users to determine their own inclusion/exclusion criteria for downstream analysis. As previewed by our applications, we foresee the Loop Catalog becoming a valuable resource for a broad range of chromatin studies including but not limited to variant-gene prioritization, machine learning, and deep learning approaches for loop prediction or utilizing loops to predict other functional measurements including gene expression as well as benchmarking analyses of such methods. At its current state, the Loop Catalog allows the community to access and bulk download uniformly processed chromatin looping data from HiChIP experiments and intermediate files, as desired. In addition, the SGL analysis, initially made available for GWAS studies in four immune-related diseases, makes it possible to query variants or genes of interest with GWAS SNPs or a prioritized subset of them such as fine-mapped SNPs. By adding SGLs derived from eQTL, we provide epigenetic context to gene regulation via genetic variation. Lastly, Loop Catalog provides 2D models of chromatin conformation overlaid with epigenetic signals to facilitate exploratory analysis and hypothesis generation in studies of 3D genome organization and gene regulation.

The Loop Catalog portal is a powerful way to access HiChIP data while providing additional analyses that are of strong interest to the scientific community. We plan to continue expanding the Loop Catalog as newly published datasets become available and our overarching, ongoing goal is to build a chromatin-centric database that seamlessly expands in terms of data as well as computational utilities offered that will eventually become the go-to platform for access and analysis of all published HiChIP data.

## Abbreviations

4DN: 4D Nucleome
AD: Atopic Dermatitis
APA: Aggregate Peak Analysis
ChIP-seq: Chromatin Immunoprecipitation-Sequencing
ChIA-PET: Chromatin Interaction Analysis with Paired-End Tag
CTCF: CCCTC-Binding Factor
dbGaP: Database of Genotypes and Phenotypes
EBI: European Bioinformatics Institute
eQTL: Expression Quantitative Trait Loci
ENCODE: ENCyclopedia Of DNA Elements
FDR: False Discovery Rate
FC Loops: FitHiChIP with ChIP-seq Peaks
FH Loops: FitHiChIP with Peaks Inferred from HiChIP
FIMO: (MEME Suite) Find Individual Motif Occurrences
GEO: Gene Expression Omnibus
GWAS: Genome-wide Association Study
HCRegLoops: High Confidence Regulatory Loops
Hi-C: High-Throughput Chromosome Conformation Capture
HiCCUPS: Hi-C Computational Unbiased Peak Search
HiChIP: Hi-C chromatin immunoprecipitation
ICE: Iterative Correction and Eigenvector Decomposition
KR: Knight and Ruiz (normalization)
LD: Linkage Disequilibrium
MAPQ: Mapping Quality (score)
NK: Natural Killer Cells
NRF: Non-Redundant Fraction
NSC: Normalized Strand Cross-Correlation Coefficient
PBC1: PCR Bottlenecking Coefficient 1
PBC2: PCR Bottlenecking Coefficient 2
QC: Quality Control
RA: Rheumatoid Arthritis
RSC: Relative Strand Cross-Correlation Coefficient
SCC: Stratum-Adjusted Correlation Coefficient
SEA: (MEME Suite) Simple Enrichment Analysis
SGL: SNP-to-Gene Linking
SNP: Single Nucleotide Polymorphism
SRA: Sequence Read Archive
PS: Psoriasis
T1D: Type 1 Diabetes
TF: Transcription Factor
TSS: Transcription Start Site
VC: Vanilla Coverage (normalization)

## Experimental design and statistics

No new experiments were performed. All data were downloaded from published data sources.

## Resources

All software and resources utilized in our work are reported in **Supp. Table S18**.

## Declarations

### Ethics approval and consent to participate

Not applicable

### Consent for publication

Not applicable

### Availability of data and materials

The Loop Catalog is freely available at https://loopcatalog.lji.org to all users without any log-in or registration requirements. The main processing pipeline has been released on Github as Loop-Catalog-Pipelines (https://github.com/ay-lab-team/Loop-Catalog-Pipelines). Similarly, we developed GEO-Resources (https://github.com/ay-lab-team/GEO-Resources) to locate HiChIP datasets in NCBI GEO, motif_pair_enrichment (https://github.com/ay-lab-team/motif_pair_enrichment) to perform enrichment analyses for motif pairs, Community-Detection-Using-Chromatin-Loops (https://github.com/ay-lab-team/Community-Detection-Using-Chromatin-Loops) to locate communities formed by chromatin loops, and T1D-Loop-Catalog (https://github.com/ay-lab-team/T1D-Loop-Catalog) to detect immune-associated SGLs. Versions of all software included in our pipeline are recorded (**Supp. Table S18**). Raw sequencing reads for HiChIP and ChIP-seq were downloaded from NCBI GEO (https://www.ncbi.nlm.nih.gov/geo/) and NCBI dbGaP (https://www.ncbi.nlm.nih.gov/gap). Hi-C contact matrices were retrieved from the 4DN Data Portal (https://data.4dnucleome.org). Fine-mapped GWAS SNPs were retrieved from CAUSALdb (http://www.mulinlab.org/causaldb). For inferring 2D embeddings for Hi-C and HiChIP data, and for their visualization, we utilized the METALoci tool from https://github.com/3DGenomes/Sox9_METALoci/tree/main

### Authors’ contributions

Joaquin Reyna: Code development, Web Development, Formal analysis, Writing—original draft.

Kyra Fetter: Data collection, Formal analysis, Writing—original draft.

Romeo Ignacio: Formal analysis, Writing—review & editing.

Cemil Can Ali Marandi: Formal analysis, Writing—review & editing.

Astoria Ma: Formal analysis. Writing—review & editing.

Nikhil Rao: Data collection, Web development.

Zichen Jiang: Data collection, Web development.

Daniela Salgado Figueroa: Data collection. Writing—review & editing.

Sourya Bhattacharya: Conceptualization, Writing—review & editing.

Ferhat Ay: Conceptualization, Writing—review & editing.

## Acknowledgements

We would like to thank Michael Talbott from La Jolla Institute for Immunology (LJI) whose web development expertise helped us set up the website and troubleshoot along the way.

## SUPPLEMENTARY FIGURE CAPTIONS

**Supp. Figure S1.** Schematic of the HiChIP and ChIP-seq data processing pipeline. Raw sequencing reads are downloaded from NCBI GEO and dbGaP and are aligned to the reference genome (hg38 or mm10). Loops are called for HiChIP (both unmerged and merged biological replicates, as indicated by the shaded circles) using HiCCUPS and FitHiChIP at the 5kb, 10kb, and 25kb resolutions. Peaks derived from both ChIP-seq and HiChIP are used for FitHiChIP loop calling. High-confidence sample sets of top H3K27ac and CTCF HiChIP samples are curated from the final set of loop calls. Red borders indicate usage in downstream database application analyses. Yellow stars indicate that the data type is available for download from the Loop Catalog web platform.

**Supp. Figure S2.** Comparison of the FitHiChIP (HiChIP-derived), HiChIP-Peaks (HiChIP-derived), and ChIPLine (ChIP-seq) peak calling methods. A) Distributions of number of peaks calls, mean peak size (bp) and median peak size (bp) for 44 human HiChIP and ChIP-seq samples from diverse cell types and protein pulldowns. B) scatter plot of recall rate versus the total peak span for peaks called by FitHiChIP (green) or HiChIP-Peaks (blue).

**Supp. Figure S3.** Reproducibility analysis for HiChIP biological replicates. A) Stratum adjusted correlation coefficients (SCCs) for all pairwise combinations of HiChIP biological replicates for 294 HiChIP samples with at least two biological replicates were generated using hicreppy. SCCs for pairwise combinations of HiChIP biological replicates are displayed for the 30 HiChIP samples with a minimum SCC below 0.90. SCCs for pairwise combinations of replicates for samples with more than 2 replicates are connected with vertical lines. Samples passed the SCC threshold if the SCC of all pairwise combinations of replicates was greater than 0.80 (n = 284 samples, a subset of 20 passing samples shown in blue). Samples which possessed at least one replicate combination with a SCC less than 0.80 did not pass the threshold (n = 10 samples, all shown in red).

**Supp. Figure S4:** Assignment of QC Flags to HIChIP Samples. Flags were assigned to all samples at the pre-processing (HiChIP and/or ChIP-seq), peak calling, and loop calling stages according to the system specified in Supp. Table S9. QC flags for FC loops consider loop quality as well as HiChIP pre-processing (PP HiChIP), ChIP-seq pre-processing (PP ChIP-seq), and ChIP-seq peak quality. Flags for FH loops are determined from loop quality, HiChIP pre-processing (HiChIP PP), and peaks inferred from HiChIP. The distribution of each flag type is displayed for A) human FC loops, unmerged and merged, B) human FH loops, unmerged and merged, C) mouse FC loops, unmerged and merged, and D) mouse FH loops, unmerged and merged.

**Supp. Figure S5.** Schema of Bootstrap Analysis for Motif Pairs. Loops are depicted with an arc and horizontal lines are used to denote their anchors. Each loop has a distinct color to emphasize the true loop composition and overlapping motifs are depicted with horizontal lines and a separate color for each (top right). Observed motif pairs are tabulated (bottom left). To simulate a new set of loops anchors are shuffled and thus bringing together new combinations of motifs (top middle). Motif pairs from these simulations are tabulated across 100000 simulations (top right) to then build a null distribution (bottom right). The observed counts for a given motif are then evaluated against their respective null distribution (bottom).

**Supp. Figure S6.** Visualization of chromatin conformation and regulatory elements surrounding genes using 2D embedding models. A) Visualization of chromatin conformation as 2D embedding models for MCM3 (visualized as a green circle). From top to bottom, the first three subpanels contain a visualization of the 2D model with bins colored based on the total number of contacts within this 4mb region (first), the level of ChIP-seq signals (second), and the local Moran’s I statistical significance value (third). The last sub panel contains a scatter plot of local Moran’s I displaying attribute (ChIP-seq signal) versus spatial lag. B-D) similar results for genes CEP85L, FAM135A, and ESR1, respectively.

**Supp. Figure S7.** Schema of the Loop Catalog Database. Tables with orange contain metadata for each HiChIP sample, purple indicates ChIP-seq data, blue indicates tables with loop or loop-associated data, and black indicates extra reference datasets (i.e. gene coordinates from GENCODE).

**Supp. Figure S8.** Selected summary of peaks and loops of the Loop Catalog. A and B) Number of peaks derived from ChIP-seq data for human and mouse, respectively. C and D) FitHiChIP loops derived for samples with ChIP-seq data at different resolutions for human and mouse, respectively.

**Supp. Figure S9.** Comparison of the HiC-Pro, distiller-nf, and Juicer for HiChIP read alignment. Scatterplots compare the number of valid pairs, cis interaction pairs, cis short-range pairs (<=20kb), cis long-range pairs (>20kb), and trans interaction pairs for A) HiC-Pro vs. distiller-nf and B) HiC-Pro vs. Juicer for 45 human HiChIP samples from diverse cell types and protein pulldowns. Distiller-nf UU (unique-unique) and UR (unique-rescue or rescue-unique) pairs were retained for this comparison, and for all tools, pairs with a cis distance less than 1kb were discarded. C) Venn diagram displaying loop overlap between loops called from pairs derived using HiC-Pro, distiller-nf, or Juicer for the CD34+-Cord-Blood.GSE165207.Homo_Sapiens.H3K27ac.b1 sample. FitHiChIP loop-calling was performed at the 5kb resolution using the stringent background model. +/− 1 bin slack was allowed. D) APA plots for all 5kb loops (top row) or the top 10,000 significant 5kb loops by q-value (bottom row) derived from pairs from each of the three alignment methods, HiC-Pro, distiller-nf, or Juicer, for the CD34+-Cord-Blood.GSE165207.Homo_Sapiens.H3K27ac.b1 sample. APA for each tool was performed using the ICE-balanced contact matrix from the same tool as background. All loops were called by FitHiChIP using ChIP-seq peaks and the stringent background model.

**Supp. Figure S10.** Comparison of the Knight-Ruiz (KR), SCALE, Vanilla Coverage (VC), and Vanilla Coverage Square Root (VC_SQRT) matrix normalization methods used for HiCCUPS loop calling. A) Distributions of number of HiCCUPS loops called at the 5kb (left), 10kb (center), and 25kb (right) resolutions from HiChIP normalized using either KR (n = 8 samples out of the 45-sample sample set for which KR converged for all chromosomes), SCALE (n=45), VC (n=45), or VC_SQRT (n=45) normalization methods. B) For the two samples H9.GSE105028.Homo_Sapiens.Rad21.b1 (top) and CD34+-Cord-Blood.GSE165207.Homo_Sapiens.H3K27ac.b1 (bottom), venn diagrams display the overlap of loops called by HiCCUPS at 5kb resolution using either SCALE, VC, or VC_SQRT normalization methods. No slack was allowed. C) For H9.GSE105028.Homo_Sapiens.Rad21.b1 (top) and CD34+-Cord-Blood.GSE165207.Homo_Sapiens.H3K27ac.b1 (bottom), APA plots display loop strength for the top significant loops by the donut FDR called by HiCCUPS using each normalization method. For H9.GSE105028.Homo_Sapiens.Rad21.b1, all samples had over 27,000 loop calls (SCALE All: n = 33,812 loops; VC All: n = 27,204 loops; VC_SQRT: n = 83,145 loops); hence, the top 10,000 significant loops were selected. For CD34+-Cord-Blood.GSE165207.Homo_Sapiens.H3K27ac.b1, VC All had 4,759 total loops (SCALE All: n = 7,126 loops; VC_SQRT: n = 18,303 loops), so 4,759 was taken as the number of top significant loops selected for each method. The ICE-normalized contact matrix generated by HiC-Pro was used as the background for all APA.

**Supp. Figure S11.** Number of FH Loop Calls by Protein Pulldown. Distributions of number of FH loop calls by protein pulldown are displayed for A) human and B) mouse samples with >100 loops. Proteins represented by >10 samples are individually displayed while all others are grouped into the “Other” category. These include proteins such as RNA-Pol-II, GATA1, STAG1, STAG2, RAD21, etc.

**Supp. Figure S12.** Loop Size of FC Loops Stratified by Protein Pulldown. Distributions of FC loop sizes for H3K27ac, CTCF and cohesin complex proteins in A) human and B) mouse samples.

**Supp. Figure S13.** Summary of SGLs derived from HiChIP and eQTL intersections. A) Breakdown by cell type for the total number of eQTLs (blue) and SNPs (orange) within the original eQTL dataset. B) Breakdown by cell type for the total number of genes from the eQTL study (green) followed by SGL summaries for unique SGLs (red), SNPs (purple) and genes (brown). * indicates a total number of SNPs or genes derived from the eQTL study before intersection with HiChIP loops.

**Supp. Figure S14.** Example analysis for CD4+ T cells using SGLs derived from their corresponding eQTLs. Depicted is the IKZF3/ORMDL3 locus with the top grey track containing SNPs derived from eQTLs followed by H3K27ac HiChIP loops for six CD4+ T cell samples.

## Supplementary Tables

**Supp. Table S1.** Summary of HiChIP Sequence Files

**Supp. Table S2.** HiC-Pro Quality Control Statistics

**Supp. Table S3.** Restriction Enzyme Sequences Used in the HiC-Pro Pipeline

**Supp. Table S4.** ChIPLine Quality Control Statistics

**Supp. Table S5.** Peak Call Statistics for HiChIP and ChIP-seq

**Supp. Table S6.** Assessment of HiChIP Reproducibility by Stratum-Adjusted Correlation Coefficients

**Supp. Table S7.** HiChIP Loop Call Statistics for FitHiChIP and HiCCUPS Loop Calling

**Supp. Table S8**: Quality Control Flags for FitHiChIP Loop Calling

**Supp. Table S9.** Quality Control Flag Definitions for FitHiChIP Loop Calling

**Supp. Table S10.** List of High Confidence HiChIP Sample Sets

**Supp. Table S11.** Hi-C Loop Call Statistics for Mustache Loop Calling

**Supp. Table S12.** GWAS SGL Statistics for RA, AD, T1D, and PS

**Supp. Table S13.** eQTL SGL Statistics for 6 Immune Cell Types

**Supp. Table S14.** 1D Conserved Anchor Motif Enrichment Analysis Statistics

**Supp. Table S15.** Paired Motif Enrichment Analysis Statistics

**Supp. Table S16.** Map of Hi-C to ChIP-seq Signals

**Supp. Table S17**. Non-Convergence of KR, SCALE, VC, and VC_SQRT Contact Matrix Normalization Methods for 45 Samples

**Supp. Table S18.** Software and Package Versions

