## Supplementary figures and images for "Loop Catalog: a comprehensive HiChIP database of human and mouse samples"

### Supp. Figure S1

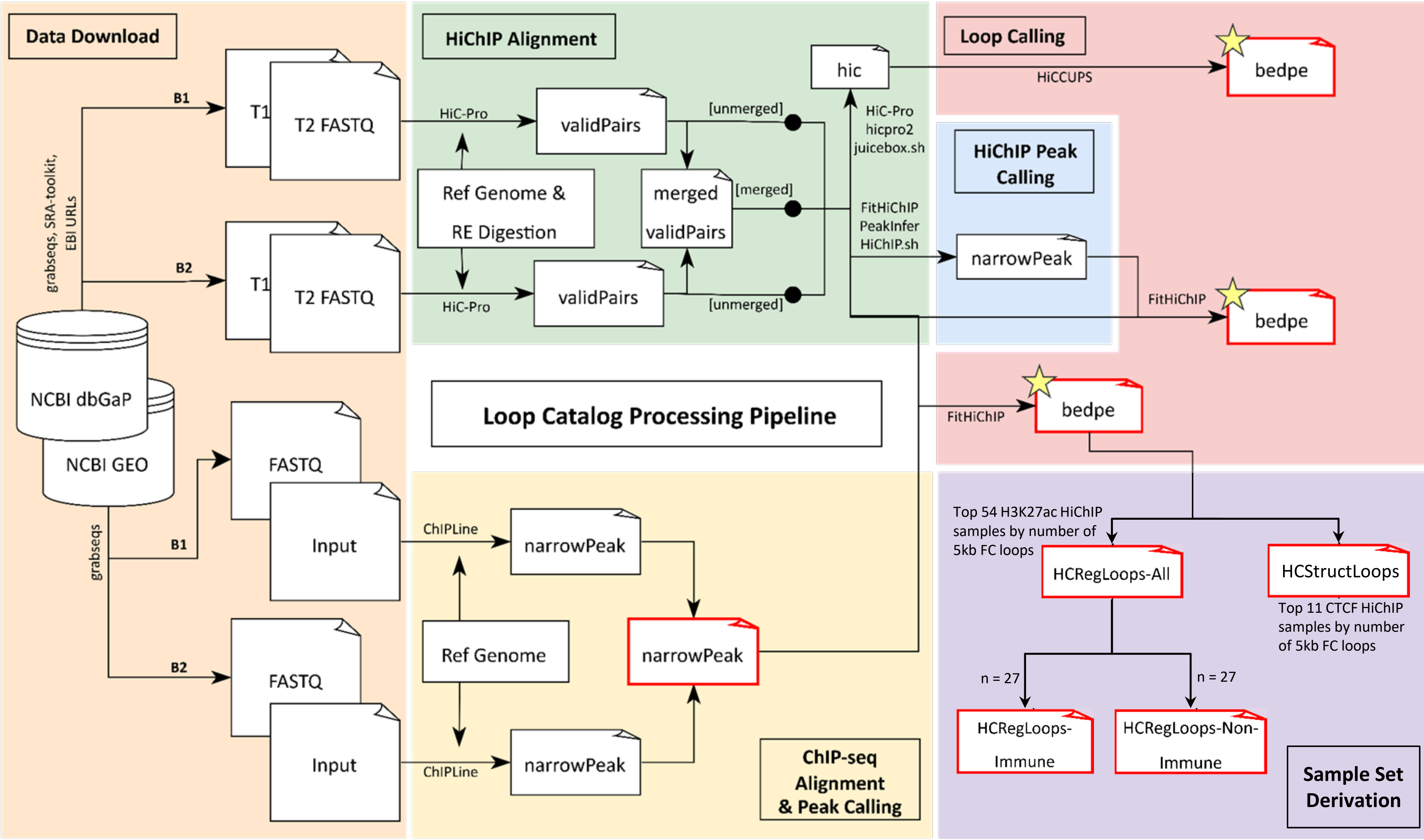

### Supp. Figure S2

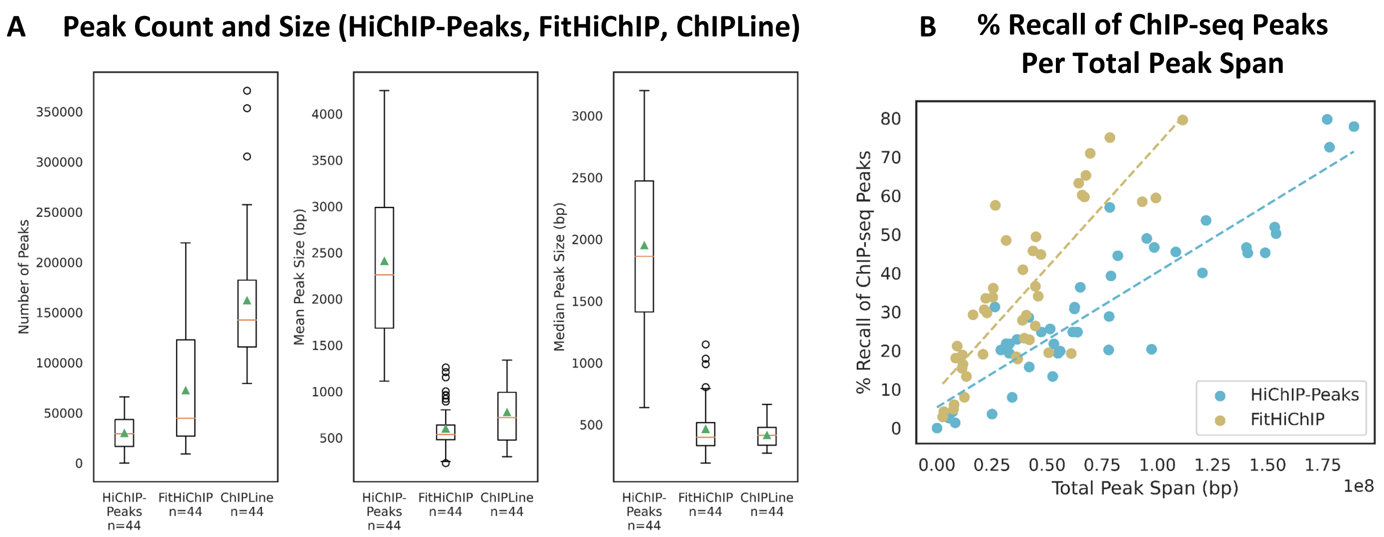

### Supp. Figure S3

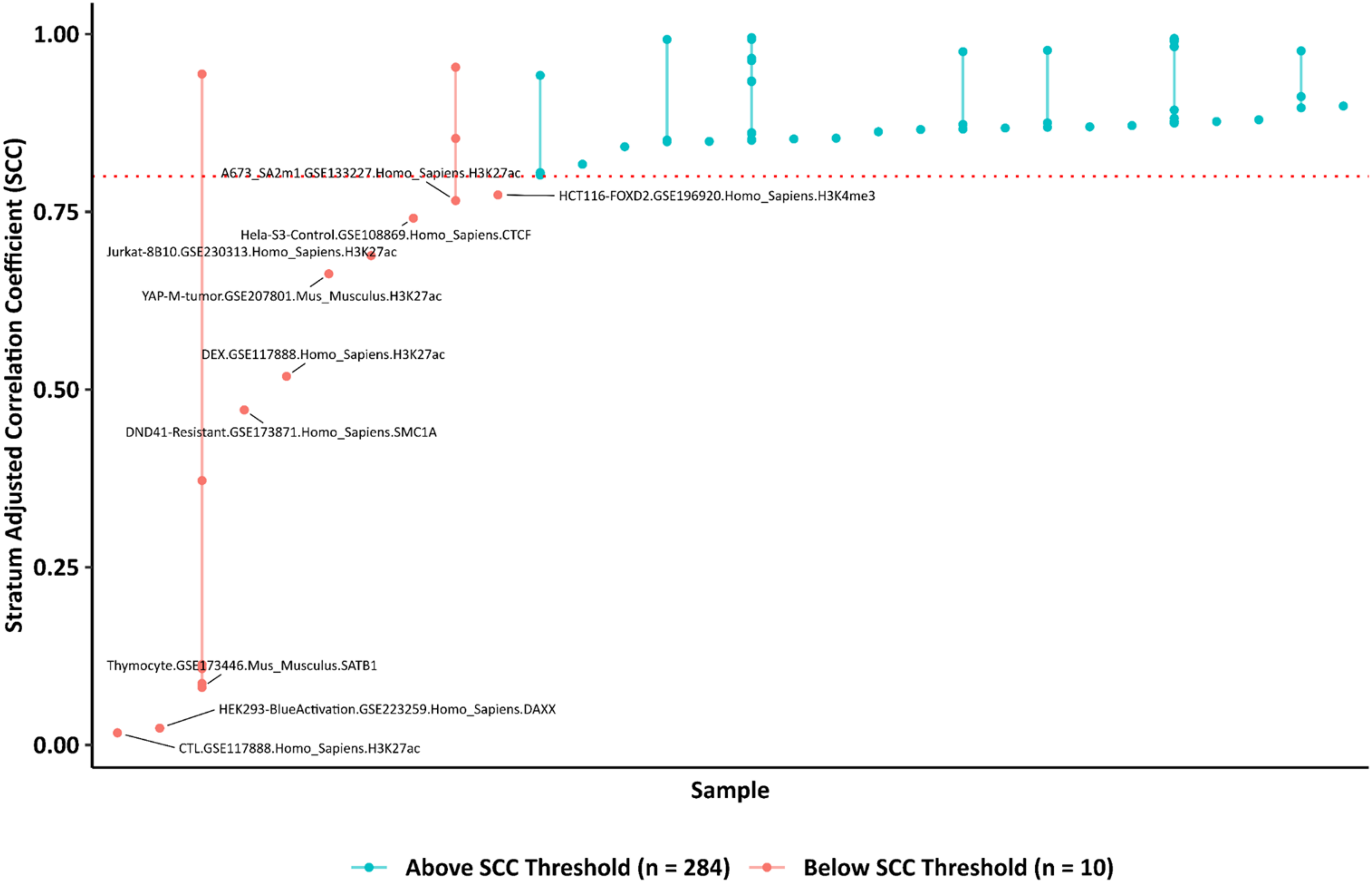

### Supp. Figure S4

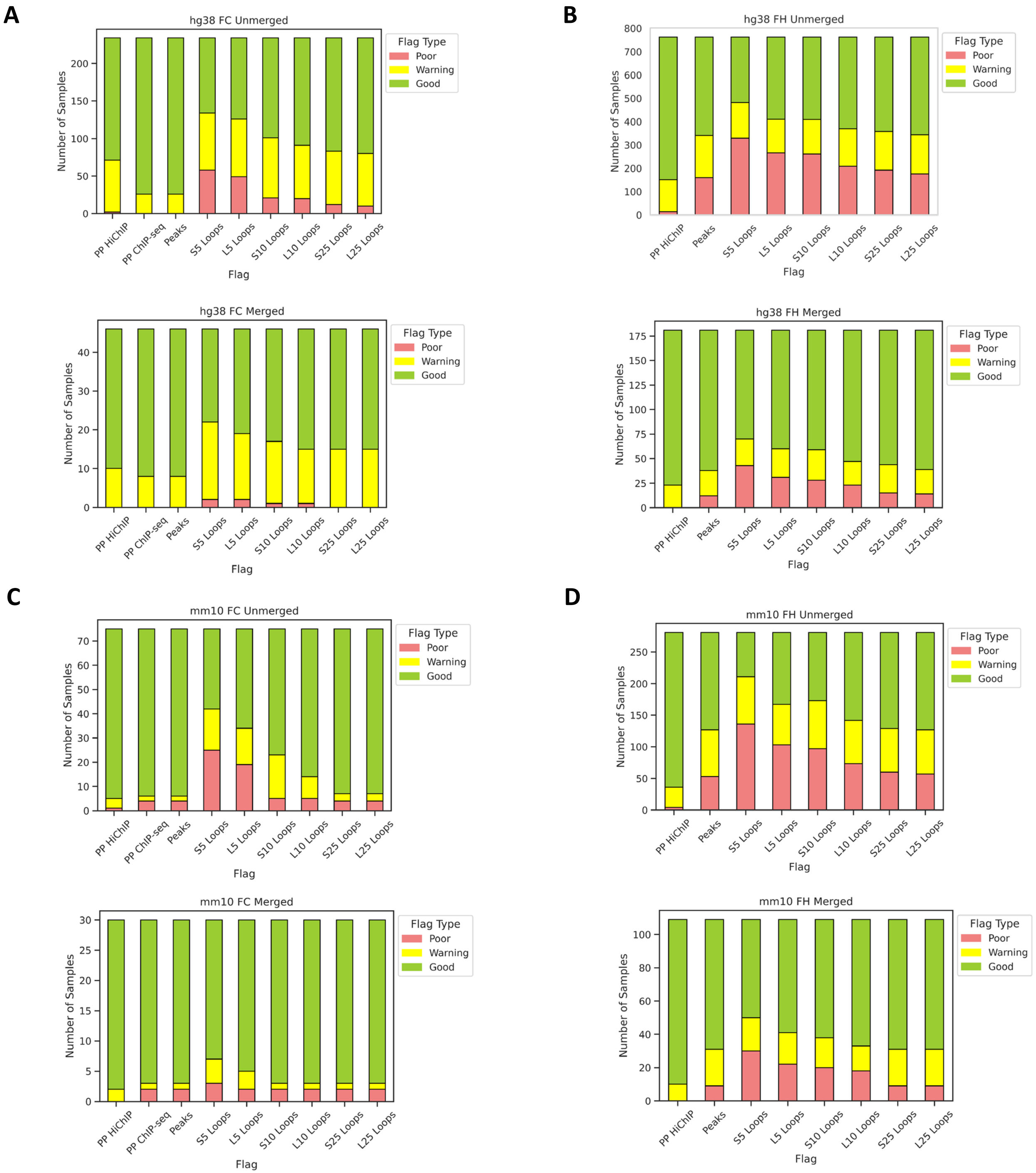

### Supp. Figure S6

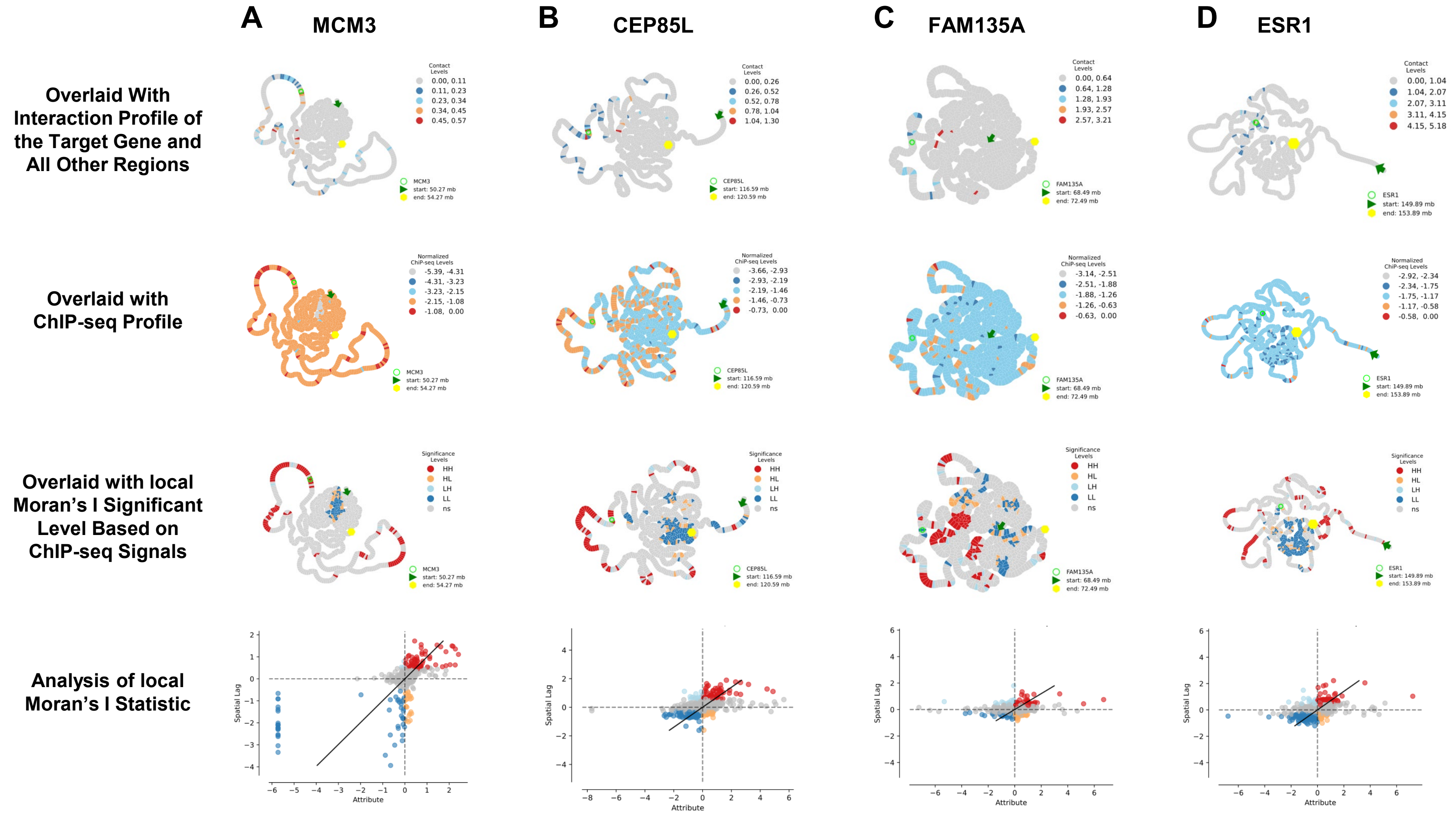

### Supp. Figure S8

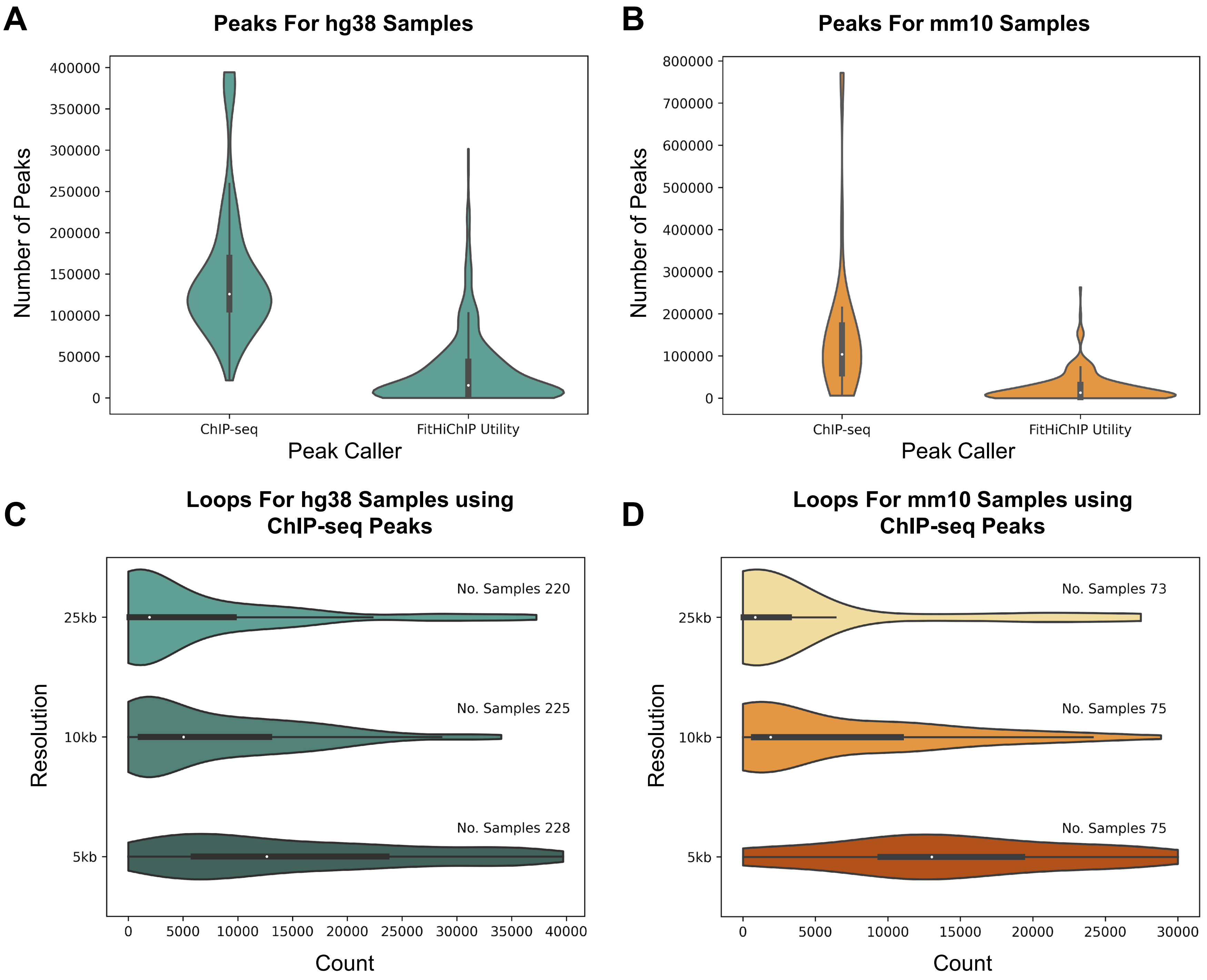

### Supp. Figure S10

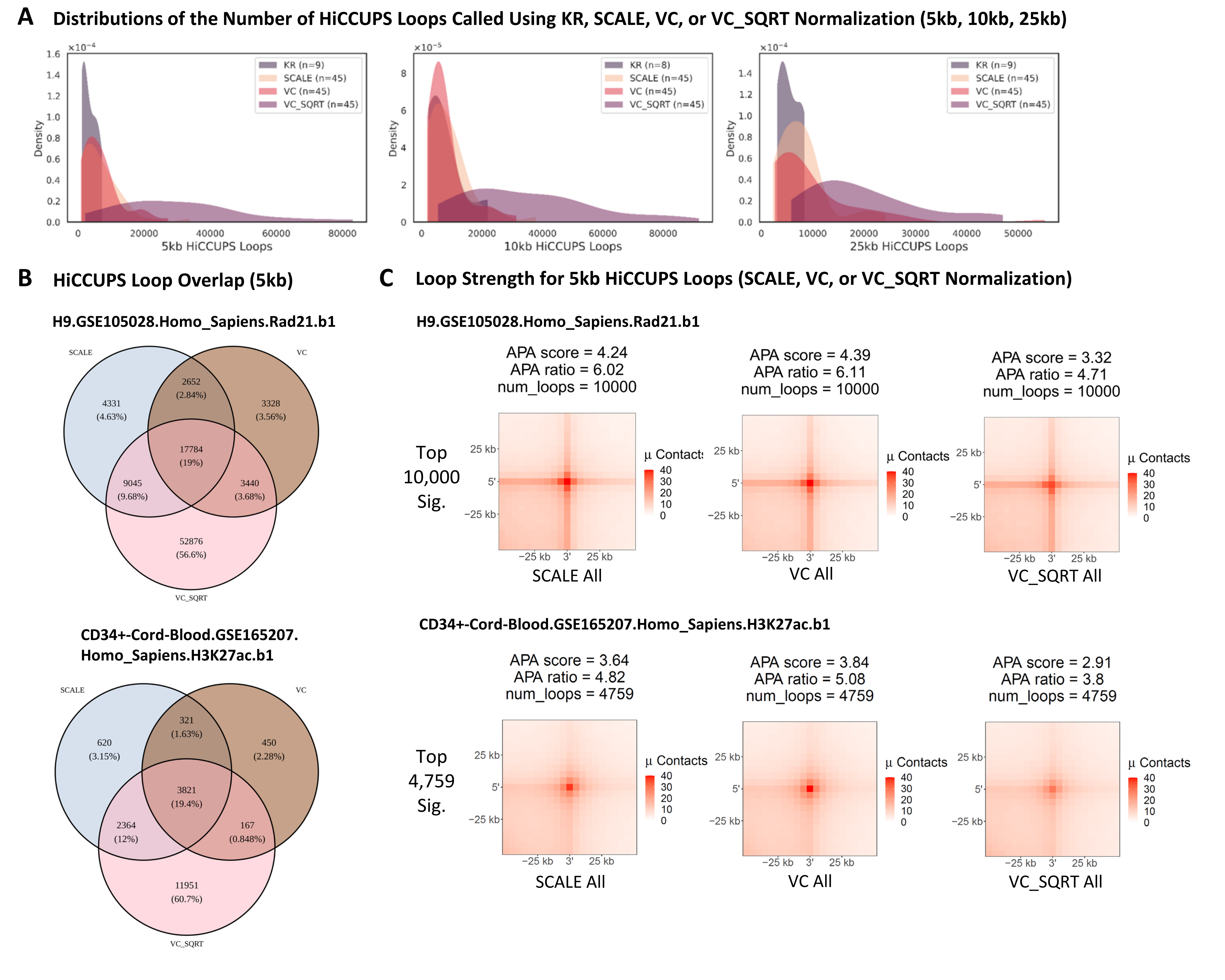

### Supp. Figure S11

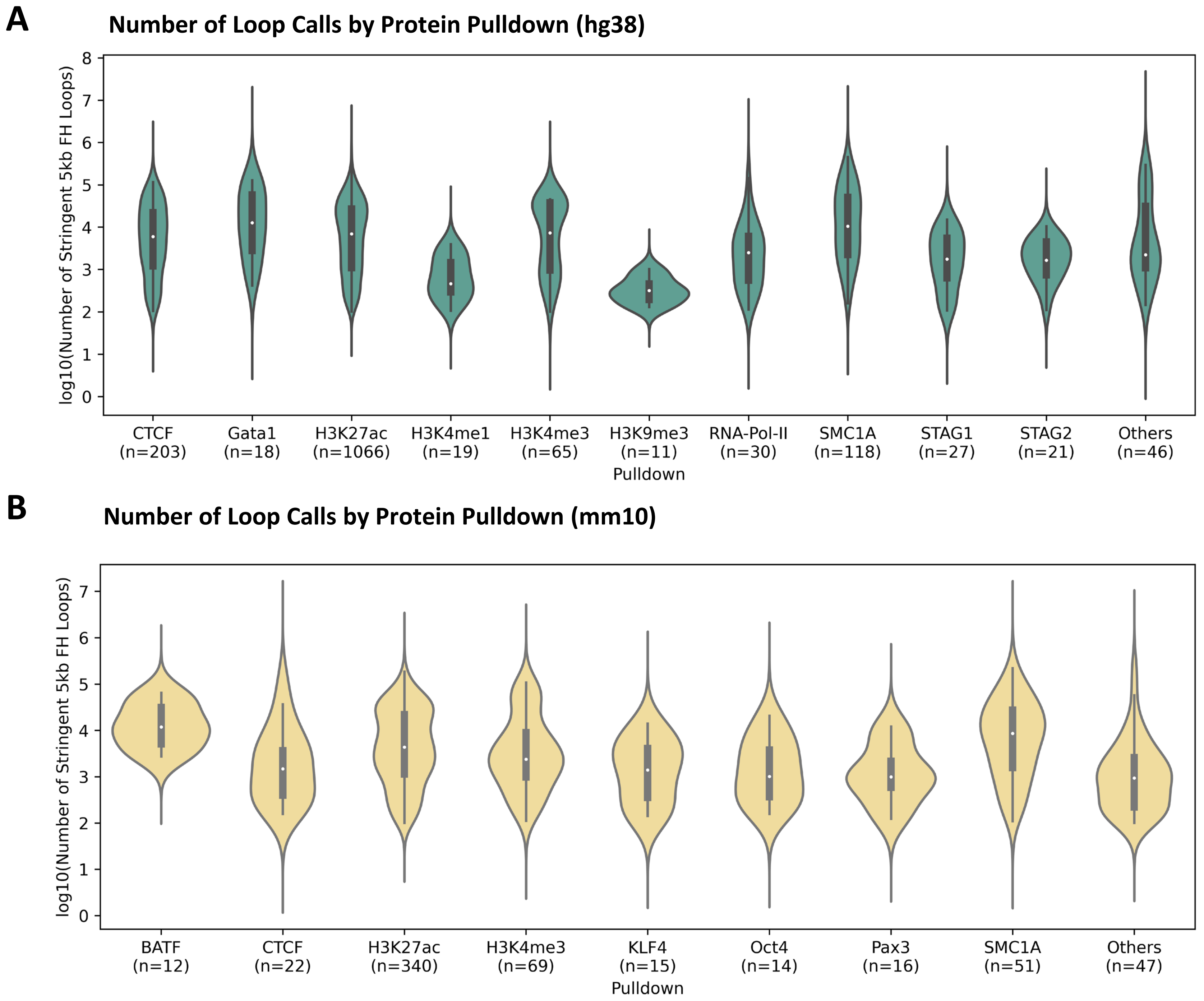

### Supp. Figure S13

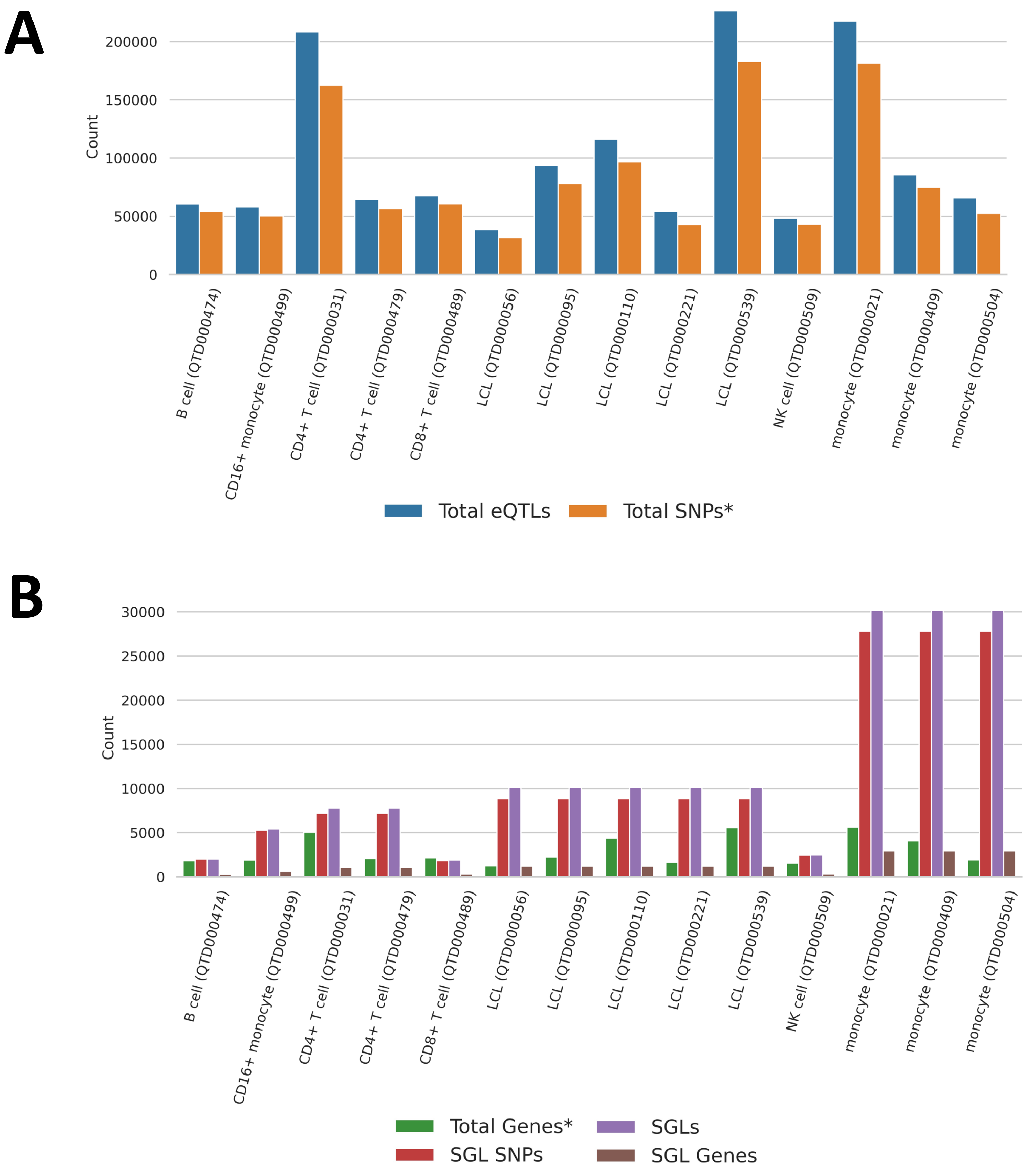

### Supp. Figure S14

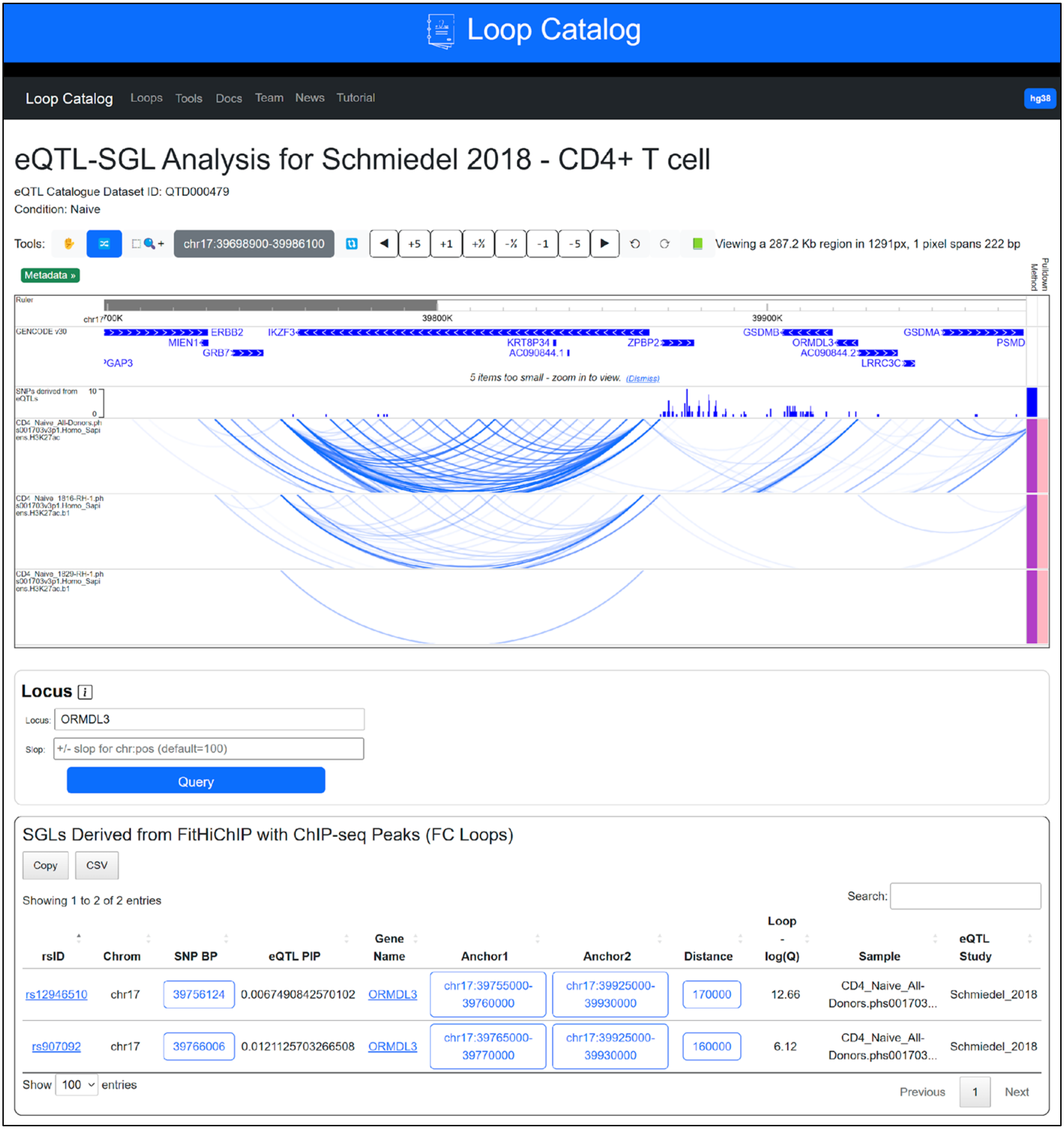
