## Supplementary material for "Loop Catalog: a comprehensive HiChIP database of human and mouse samples": Supp. Figure S5

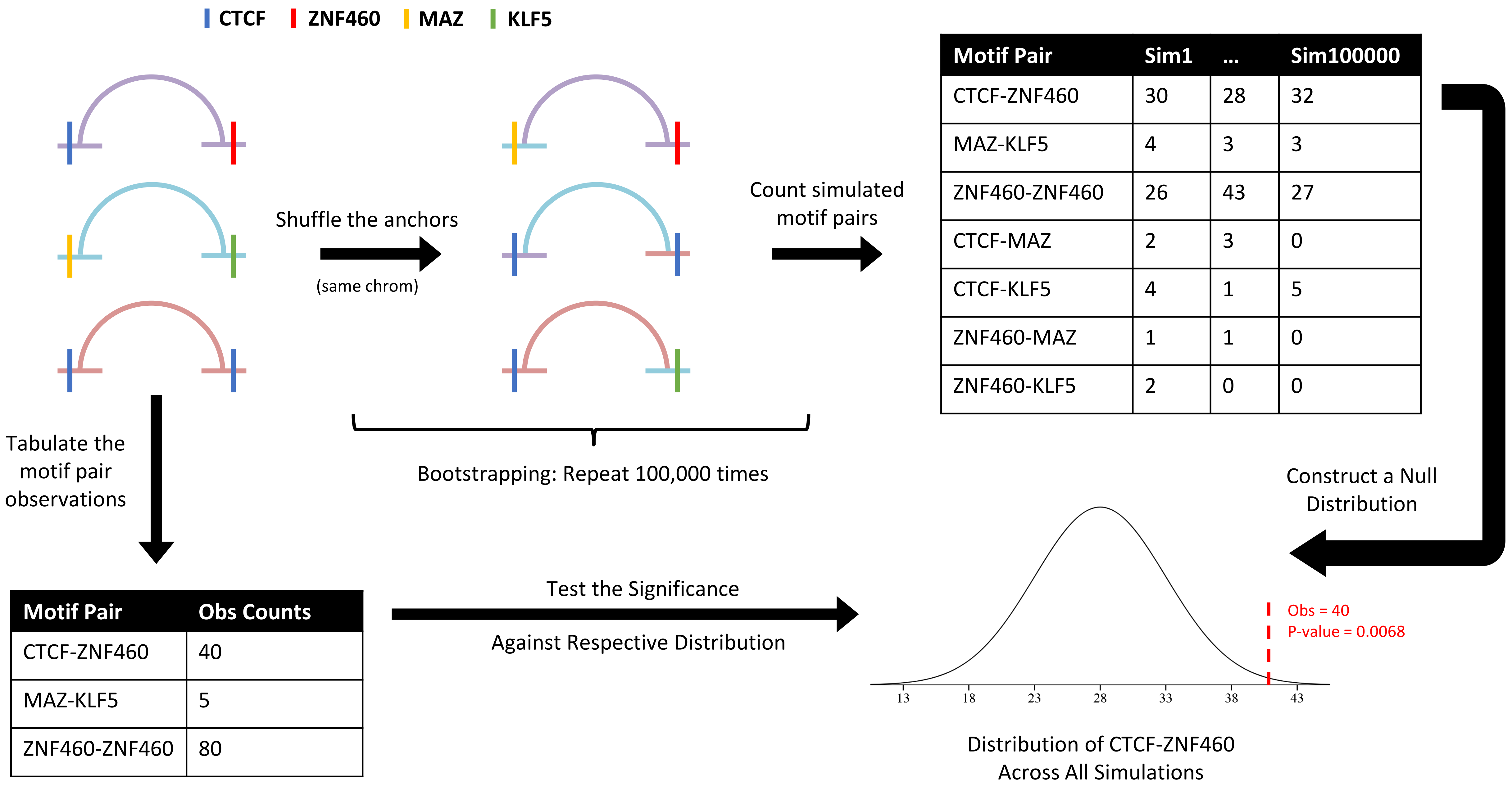


| Motif Pair | Sim1 | ... | Sim100000 |
|---|---|---|---|
| CTCF-ZNF460 | 30 | 28 | 32 |
| MAZ-KLF5 | 4 | 3 | 3 |
| ZNF460-ZNF460 | 26 | 43 | 27 |
| CTCF-MAZ | 2 | 3 | 0 |
| CTCF-KLF5 | 4 | 1 | 5 |
| ZNF460-MAZ | 1 | 1 | 0 |
| ZNF460-KLF5 | 2 | 0 | 0 |

| Motif Pair | Obs Counts |
|---|---|
| CTCF-ZNF460 | 40 |
| MAZ-KLF5 | 5 |
| ZNF460-ZNF460 | 80 |
