## Supplementary material for "Loop Catalog: a comprehensive HiChIP database of human and mouse samples": Supp. Figure S7

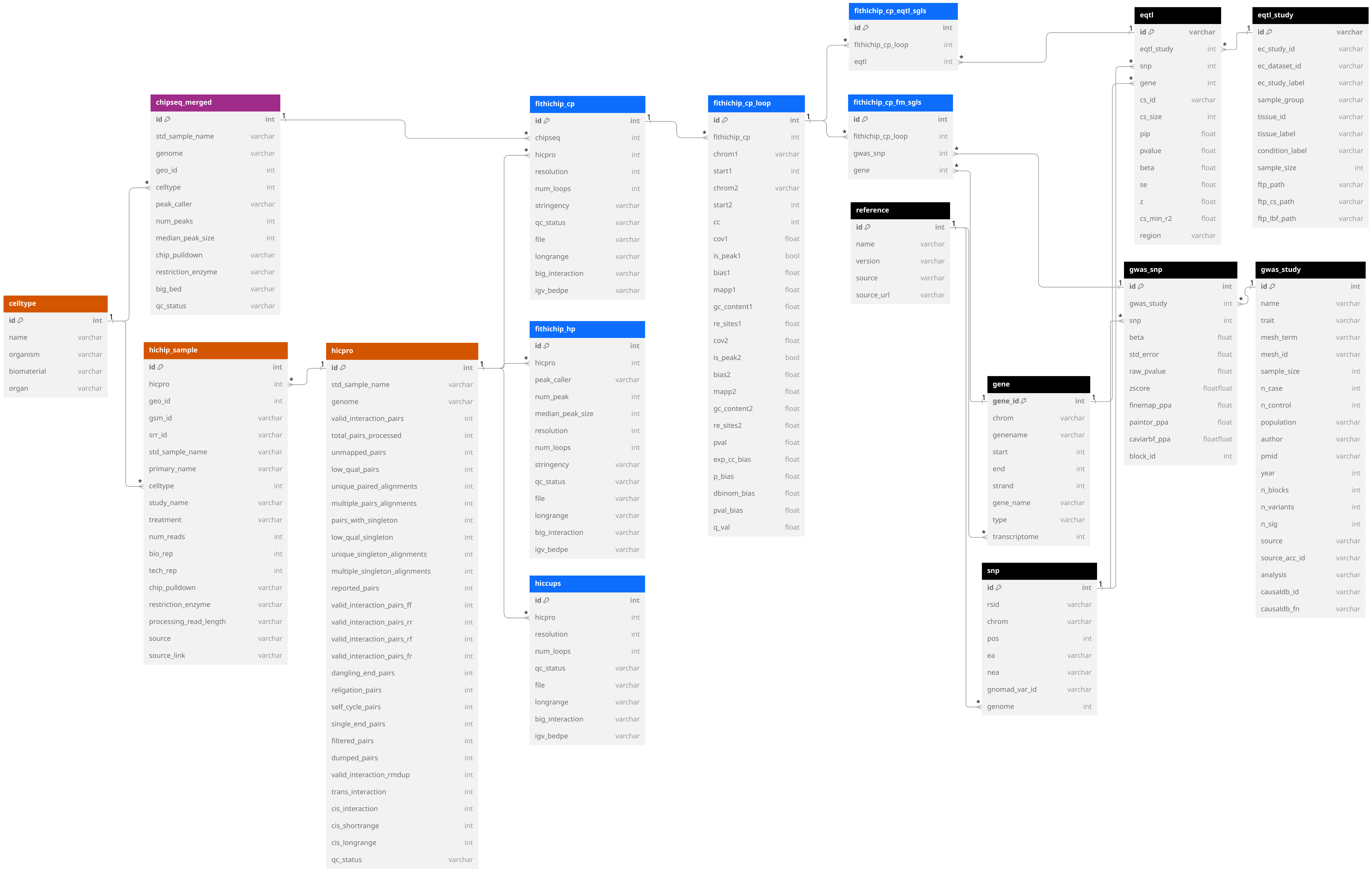


Supplementary Figure 7. Schema of the Loop Catalog Database. Tables with orange contain metadata for each HiChIP sample, purple indicates ChIP-seq data, blue indicates tables with loop or loop-associated data, and black indicates extra reference datasets (i.e. gene coordinates from GENCODE).
