## Supplementary material for "Loop Catalog: a comprehensive HiChIP database of human and mouse samples": Supp. Figure S9

**A Comparison of Alignment Metrics (HiC-Pro vs. distiller-nf)**

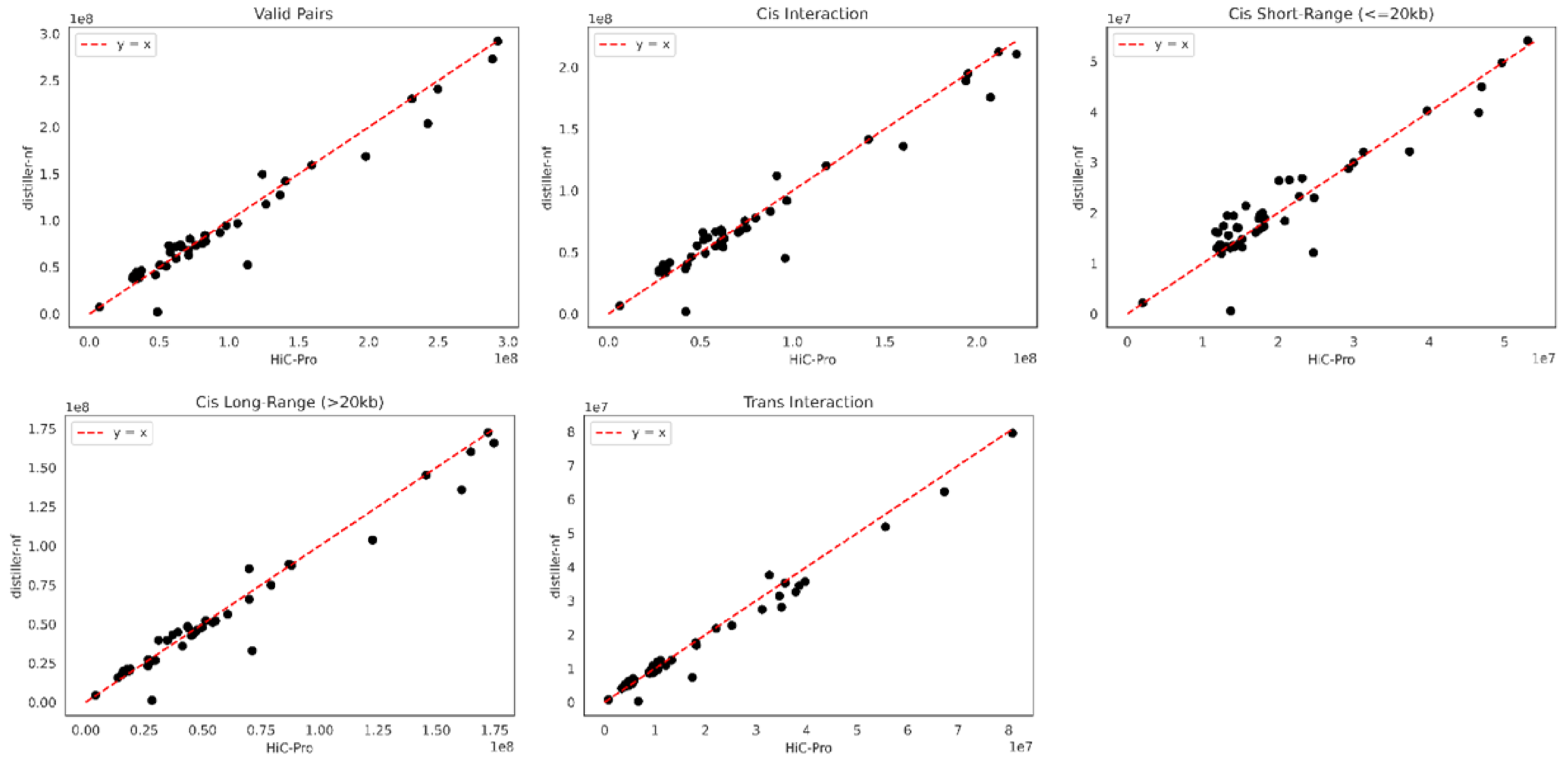


**B Comparison of Alignment Metrics (HiC-Pro vs. Juicer)**

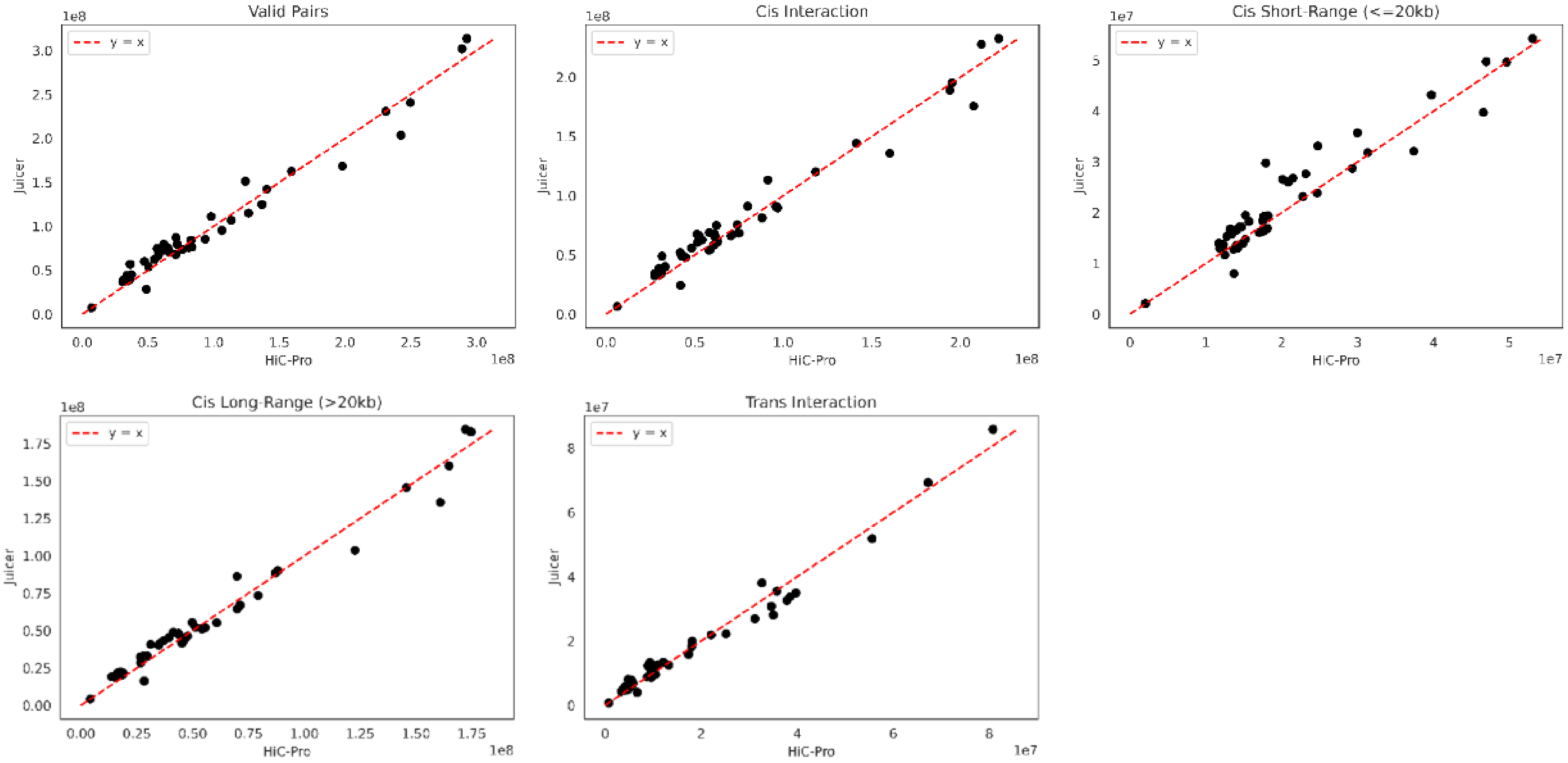


**C Loop Overlap (1kb Slack)**

CD34+-Cord-Blood.GSE165207.Homo_Sapiens.H3K27ac.b1

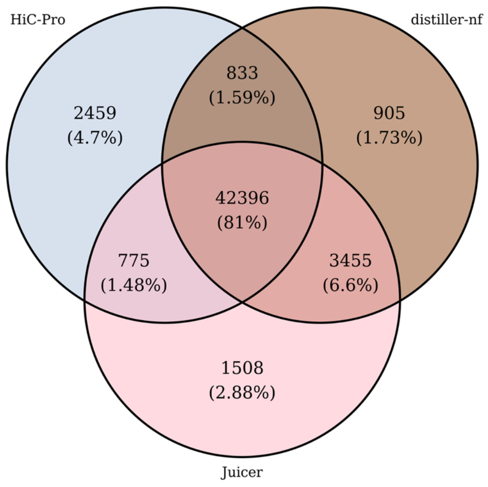


**D Loop Strength for Loops Derived from HiC-Pro, distiller-nf, or Juicer**

CD34+-Cord-Blood.GSE165207.Homo_Sapiens.H3K27ac.b1

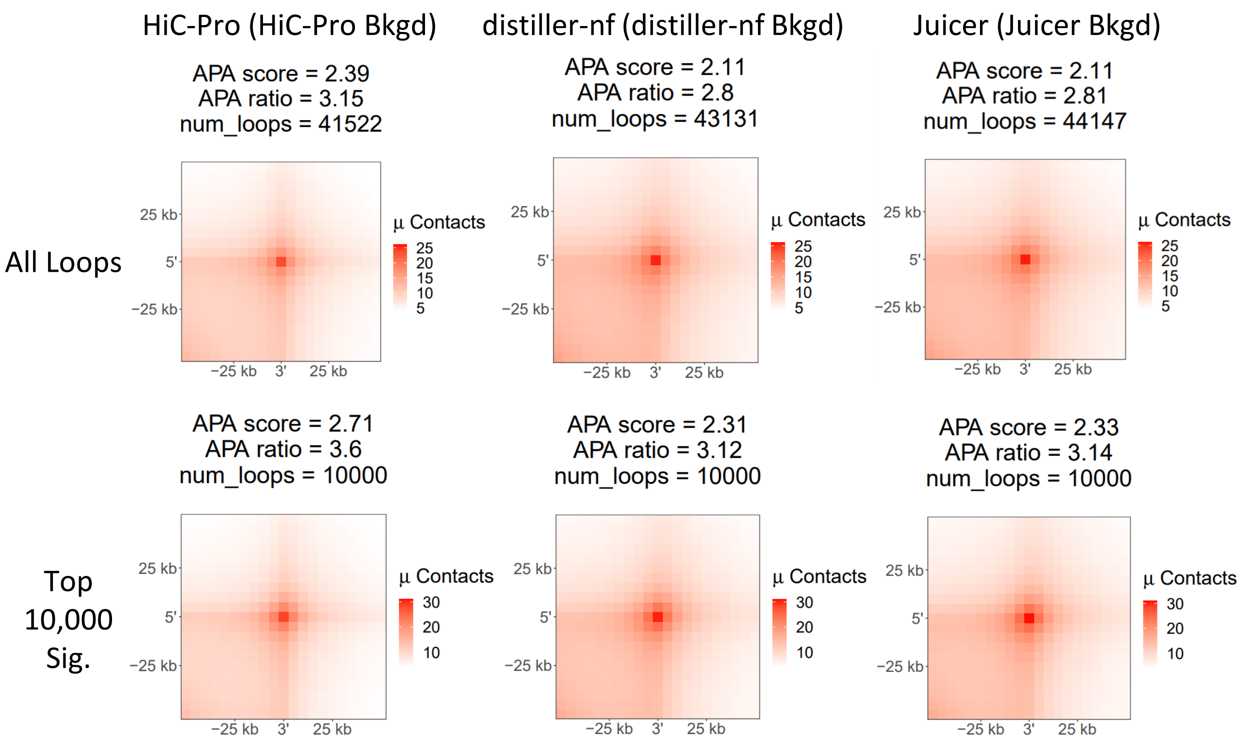
