## Supplementary material for "Loop Catalog: a comprehensive HiChIP database of human and mouse samples": Supp. Figure S12

**A** **Distribution of Loops Sizes for 5kb FC Loops (distance < 1 million bp) (hg38)**

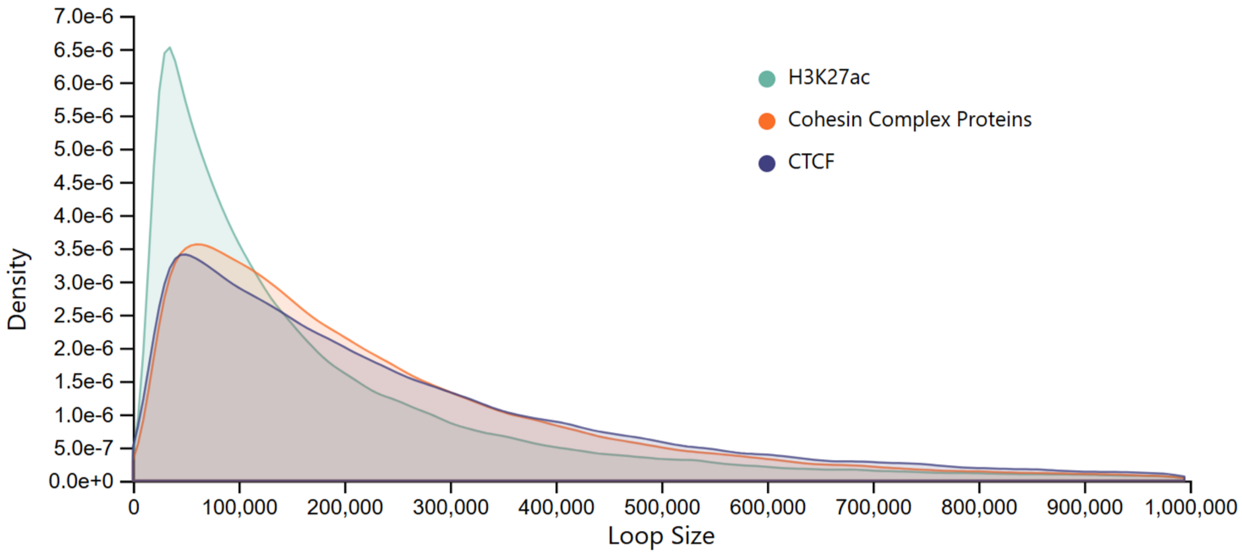


**B** **Distribution of Loops Sizes for 5kb FC Loops (distance < 1 million bp) (mm10)**

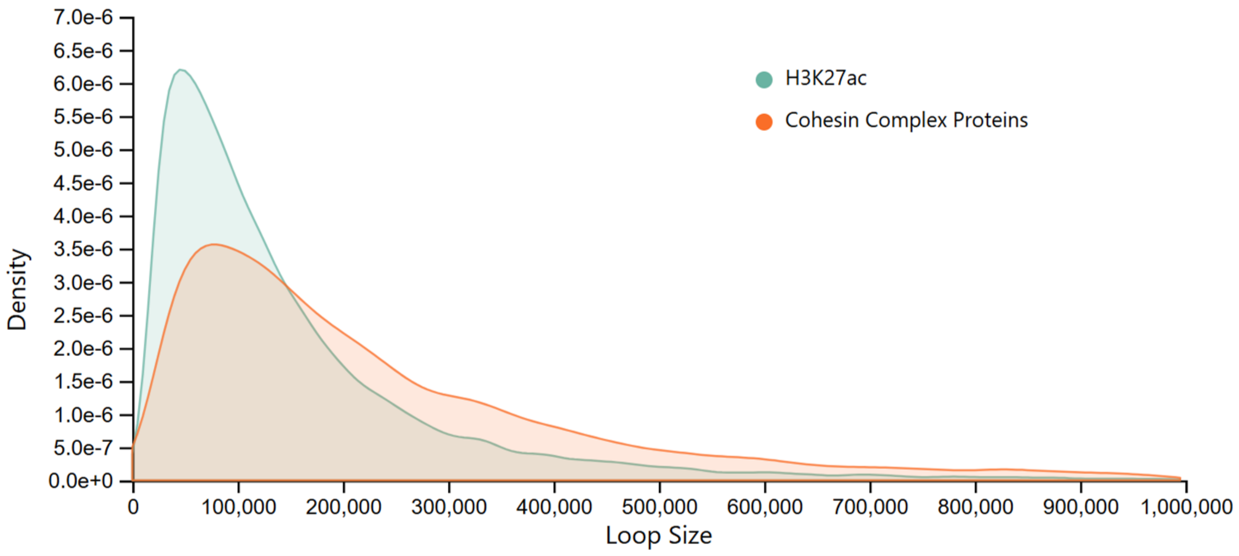
